# The carbon-nitrogen metabolism gene E1OGDH1 influences maize root system architecture and nitrogen plasticity

**DOI:** 10.64898/2026.07.21.739812

**Authors:** Michelle S. Cho, Zhengbin Liu, Collin Luebbert, Greg Ziegler, Dhineshkumar Thiruppathi, Christine Tiskevich, Hua Liu, Bing Yang, Ivan Baxter, Stephen Moose, Christopher N. Topp

**Affiliations:** Division of Biology and Biomedical Studies, Department of Biology, Washington University in St. Louis, St. Louis, MO, 63110 USA; Donald Danforth Plant Science Center, St. Louis, MO, 63132 USA; Department of Crop Sciences, University of Illinois Urbana-Champaign, Urbana, IL 61801, USA; Division of Plant Science and Technology, Bond Life Sciences Center, University of Missouri, Columbia, MO 65211, USA; Michigan State University, Plant, Soil and Microbial Sciences and Plant Biology: Plant and Soil Sciences Building 1066 Bogue Street, East Lansing, MI 48824, USA

## Abstract

Synthetic nitrogen fertilizers have greatly increased crop yields, yet much of the applied nitrogen is lost from agroecosystems and contributes to environmental pollution and higher economic costs. Improving nitrogen uptake efficiency (NUpE) benefits from understanding how root system architecture (RSA) governs soil nitrogen capture. Although root traits have seldom been explicit breeding targets, selection for variation in above-ground nitrogen accumulation has also likely shaped differences in RSA. The Illinois Protein Strain Recombinant Inbred population, derived from more than a century of divergent selection for seed protein concentration, offers a powerful resource for dissecting RSA variation. Using multi-year field phenotyping of excavated root crowns and genome-wide association analysis, we identified a quantitative trait locus on chromosome 10 containing E1OGDH1, which encodes the E1 subunit of the 2-oxoglutarate dehydrogenase (OGDH) complex. OGDH performs a key step in the tricarboxylic acid cycle that also modulates 2-oxoglutarate, an important entry point into nitrogen metabolism and a co-factor for enzymes involved in hormone and secondary product synthesis. Long-read sequencing of inbreds derived from the divergent IHP and ILP parental populations revealed promoter polymorphisms defining E1OGDH1 alleles and differed in E1OGDH1 expression in root tissue. Field experiments in IPSRI lines carrying IHP- or ILP-associated E1OGDH1 alleles showed differences in root architectural traits over two years. CRISPR-Cas9 knockout mutants confirmed a functional role for E1OGDH1 in whole-plant performance and nitrogen-responsive root development. Mutants were shorter, had reduced biomass, and exhibited altered architectural responses to soil nitrogen levels. Transcriptome analysis further showed that loss of E1OGDH1 altered basal and nitrogen-responsive expression of genes associated with root development and nitrogen uptake and metabolism. Together, these findings identify E1OGDH1 as a strong candidate quantitative regulator of maize RSA and nitrogen plasticity, suggesting that central carbon–nitrogen metabolic genes can contribute to root developmental responses relevant to NUpE.

## Introduction

Nitrogen Use Efficiency (NUE) is a critical trait for plant breeding due to its significant environmental and economic implications. The Green Revolution led to substantial increases in crop yields without increasing agricultural land use through multiple tactics such as the development of high-yielding varieties, mechanization, and the increased use of chemical nitrogen fertilizers (Mann, 1997). Maize breeding in the US has increased the NUE and harvest index of modern hybrids (Mueller, Messina and Vyn, 2019; Ruiz *et al*., 2023), but they still typically capture only 50% or less of applied nitrogen (Raun and Johnson, 1999; Zhu and Chen, 2002). Soil, air, and water pollution occur during the production, transportation, application, and runoff of nitrogen fertilizers. This inefficiency in nitrogen uptake poses ecological challenges and presents economic constraints, particularly for developing nations that lack the means to deploy nitrogen fertilizers and close the “yield gap” relative to optimal production (Mann, 1997; Tilman *et al*., 2002; Good and Beatty, 2011; West *et al*., 2014; Pradhan *et al*., 2015). Therefore, understanding nitrogen metabolism in crops and, ultimately, high NUE crops has become a desirable target for breeding.

Variation in root system architecture (RSA), the spatiotemporal deployment of the root system, determines the efficacy and efficiency of plant nutrient uptake, yet is relatively understudied compared to above-ground traits. Since nitrogen is a major driver of growth, plants are tuned to both internal physiological and external soil nitrogen status. While most genotypes develop steeper roots under nitrogen deficiency to access deeper nitrogen pools, significant genotypic variation exists in the direction and magnitude of this plastic response, with a subset of genotypes showing shallower or unchanged root angles under nitrogen limitation (Trachsel *et al*., 2013). In a study observing teosinte, the origin of the modern corn, and the W22 inbred line in aeroponic conditions, crown root numbers were seen to decrease, while their lengths increased in low nitrogen conditions (Gaudin *et al*., 2011; Gaudin, McClymont and Raizada, 2011). Similarly, low nitrogen stress in hydroponic conditions increased axile root elongation, reduced the number of crown roots, and decreased lateral root density and length in maize (Gao *et al*., 2015). Lateral root branching density is influenced by the availability of nitrogen and phosphorus, with nitrogen being more mobile than phosphorus in soil(Postma, Dathe and Lynch, 2014). Nitrogen stress in greenhouse-grown maize led to node-specific root system responses, with fewer developed nodal roots, but increased occupancy in higher nodes (Schneider *et al*., 2021). In other crops and environments, such as rapeseed (*Brassica napus* L.) in hydroponically grown conditions, low nitrate increases the root-to-shoot biomass ratio and promotes primary root growth, with lateral root responses varying by population (Haelterman *et al*., 2025). In rice in low nitrogen soil pot conditions, root length, surface area, root number, root volume, and root cortical area decreased (Liu *et al*., 2023). Together, these findings highlight that nitrogen limitation drives genotype- and environment-specific changes in root system architecture, though the direction and functional consequences of these changes for nitrogen acquisition remain context-dependent and are not yet fully resolved. Establishing clearer links between RSA traits and nitrogen uptake capacity in mature, field-grown plants and using sufficiently powered genetic populations therefore remains a critical and unmet challenge in root biology.

A powerful resource to explore genetic variation in NUE is the Illinois Long Term Selection Experiment (ILTSE), where more than 100 generations of divergent recurrent selection have increased or decreased kernel protein and thus nitrogen storage in seeds. This experiment began in 1896 with an open-pollinated variety of maize, Burr’s White. The resulting populations, Illinois High Protein (IHP, 30% protein) and Illinois Low Protein (ILP, 4% protein), have shifted approximately threefold in both directions from the original population mean (Moose, Dudley and Rocheford, 2004). Using the 70th generation of the Illinois High Protein (IHP) and Illinois Low Protein (ILP) populations in a multiparent crossing scheme, 500 Illinois Protein Strain Recombinant Inbreds (IPSRIs) were made (Dudley, 2007; Lucas *et al*., 2013). This scheme included seven generations of random mating to reduce linkage disequilibrium prior to inbreeding. Using this intermated Recombinant Inbred Line (RIL) population, molecular marker studies are possible (Goldman, Rocheford and Dudley, 1993, 1994; Alrefai, Orozco and Rocheford, 1994; Mikkilineni and Rocheford, 2004; Dudley, 2007). Prior work (Uribelarrea, Below and Moose, 2004) comparing the two inbreds, IHP1 and ILP1, derived from cycle 90 of each population, showed that uptake efficiency and total plant nitrogen accumulation are significantly greater in IHP1 than in ILP1 across nitrogen levels tested in the field (Uribelarrea, Below and Moose, 2004; Uribelarrea, Crafts-Brandner and Below, 2008). Seed protein concentration also varies widely in the IPSRIs (from 10-21%, Lucas et al., 2013).

Because the nitrogen accumulated in seed ultimately depends on nitrogen acquired by the root system, we reasoned that long-term divergence in kernel nitrogen demand could have indirectly affected root traits, root-associated physiology, or linked genetic loci influencing nitrogen acquisition. We therefore did not assume that selection for high or low seed protein directly caused specific RSA phenotypes. Rather, we hypothesized that the IPSRI population, would segregate for genetic variation affecting root system architecture. Consistent with this expectation, we subsequently demonstrated that 3D root system architectural differences among IHP1, ILP1, and the IPSRIs are evident in 3-week-old plants and identified quantitative trait loci controlling these traits (Li et al., 2023).

However, identifying causal genes from GWAS in mature root systems grown in the field remains a major challenge due to the limitations of field root phenotyping, including environmental influence on RSA (Topp *et al*., 2016). Their underground nature makes real-time, non-destructive measurements difficult, and thus only a scant few examples have been published in any cereal crop (Kirschner *et al*., 2024) including maize (Schneider *et al*., 2021; Ren *et al*., 2022). Yet recent advances in image-based phenomics that increase the accuracy and amount of measurements have demonstrated that phenotype-to-genotype associations can be improved (M. Li *et al*., 2018; Liu *et al*., 2021; Shao *et al*., 2021; Li *et al*., 2023; Hein *et al*., 2025). To identify new genetic regulators of root system architecture that may increase nitrogen uptake efficiency, we conducted a multi-year field experiment, applying cutting-edge 2D and 3D root phenomics technologies to the IPSRI recombinant inbred lines. GWAS on dozens of phenotypes extracted from digital images of excavated root crowns identified several traits associated with single-nucleotide polymorphisms (SNPs), referred to here as trait-associated SNPs (TAS).

We focused on a region of TAS on chromosome 10 that included a gene encoding the 2-oxoglutarate dehydrogenase E1 subunit. This protein is one part of the multimeric protein complex 2-oxoglutarate dehydrogenase (OGDH) complex, which in plants is a crucial enzyme in the TCA cycle, catalyzing the oxidative decarboxylation of 2-oxoglutarate to succinyl-CoA. The complex consists of three main components: E1 (2-oxoglutarate dehydrogenase), E2 (dihydrolipoyl succinyltransferase), and E3 (dihydrolipoamide dehydrogenase) (Miller *et al*., 2008). The E1 subunit initiates the reaction by decarboxylating 2-oxoglutarate, the E2 subunit transfers the succinyl group to CoA, and the E3 subunit regenerates the oxidized form of lipoamide (Frank *et al*., 2007). The substrate 2-oxoglutarate can be used in the TCA cycle for mitochondrial respiration, or it may be converted to glutamate for assimilation of nitrate and transamination into other amino acids. Therefore, this process is essential for plant metabolism, in carbon-nitrogen interactions, and is considered a limiting step in the TCA cycle. The OGDH complex’s activity is regulated through allosteric mechanisms and reversible phosphorylation, allowing plants to fine-tune TCA cycle flux in response to metabolic needs (Frank *et al*., 2007). Studies on Arabidopsis mutants have shown that disruption of OGDH complex components can lead to altered respiration rates, changes in carbohydrate metabolism, smaller roots, impacts on plant growth, and seed production (Millar, Hill and Leaver, 1999; Condori-Apfata *et al*., 2019).

Sequencing of E1OGDH1 in IHP1 and ILP1, inbred lines derived from the IHP and ILP parental populations, revealed promoter polymorphisms which were associated with differential expression levels. Further field experiments showed consistent differences across years for root phenotypes among IPSRIs carrying IHP1 compared to ILP1 alleles at E1OGDH1. Functional validation through E1OGDH1 knockout mutants demonstrated reduced plant height, biomass, and altered RSA. Notably, several root traits and numerous genes influencing root development, nitrogen import, and nitrogen metabolism differed between mutant and wild type and/or were responsive to nitrogen availability, and we observed significant interactions between genotype and nitrogen treatment. Together, these findings support a functional role for E1OGDH1 in regulating root architecture and nitrogen responsiveness, highlighting it as a promising genetic target for improving nitrogen use efficiency in maize.

## Results

### GWAS using digital root traits of the field excavated root identified numerous Trait Associated SNPs

The IHP1 and ILP1 inbreds derived from the ILTSE experiment (Uribelarrea, Below and Moose, 2004) vary dramatically for total plant nitrogen accumulation and seed protein concentration, with IHP1 showing increased nitrogen amounts relative to ILP1 at all the nitrogen levels tested in the field (Uribelarrea, Crafts-Brandner and Below, 2008). We reasoned that changes in RSA might support the higher N uptake of IHP1, and confirmed nearly twice as many seedling lateral roots in IHP1 compared to ILP1 imaged in a 3-D gel system (Li *et al*., 2023). These encouraging results led us to evaluate variation within the Illinois Protein Selection Recombinant Inbred (IPSRI) mapping population grown in the field. To make feasible the multi-year root phenotyping campaign, we selected 138 “core” lines, a subset of the total 500-line IPSRI population. The core IPSRIs retain the limited population structure and distribution of kernel nitrogen concentration present in the full 500, based on prior genotyping at selected loci (Dudley, 2007; Lucas *et al*., 2013; Supplementary Figure 1). The maize plants were grown until maturity and the roots were methodically excavated with a shovel (Fig. 1A). Root samples were washed, imaged by 2D photography, and the images were analyzed using Digital Imaging of Root Traits (DIRT), enabling rapid, reliable, and increased feature extraction compared to manual measurements (Das *et al*., 2015). 53 root architecture traits from the IPSRI population were used to perform a Genome-Wide Association Study (GWAS) (Methods) (Fig. 1). The broad-sense heritability of the root architecture traits ranged from 24.8% to 0%. Grain protein concentration was also measured in 2014 from ears harvested from the same plants sampled for roots. Grain protein concentration varied from 7.5 – 20.7% among the IPSRIs, but was only weakly correlated with four DIRT pipeline traits: RTA dominant angle (0.20), CPD 25 (-0.15), STA median (0.14) and average lateral length (-0.14). Single-nucleotide polymorphisms (SNPs) were filtered for minor allele frequency above 5% and missing rate below 10%, retaining 95,680 markers. Missing calls were imputed to the mean allele at that SNP, and principal components (PC’s) of the snp matrix were calculated. A generalized linear model with the first 5 PC’s fit as covariates was used to perform genome-wide association studies (GWAS) for 80 Multivariate Adaptive Shrinkage (MASH) analyses and 87 root crown traits in 2014 and 2015, respectively. A significance threshold was established by first identifying 7,052 independent SNP’s by clustering by linkage disequilibrium (LD) using a threshold of R² > 0.2. This set of independent SNP’s was then used to establish a less conservative Bonferroni correction. 19 traits in 2014 and 23 traits in 2015 displayed ρ-values below this adjusted significance. To identify TAS that had statistical support across multiple years and traits, we employed a Bayesian learning model that computes posterior effect estimates and significance scores based on GWAS summary statistics (Urbut *et al*., 2019) and found 1,361 SNPs with a log10 Bayes factor > 5 (Fig. 1B). Candidate windows around mash hits were created by considering SNPs in LD with a given mash-identified SNP at R² > 0.25. Therefore, by using the IPSRI population root traits, we were able to find several root trait-associated SNPs that could contribute to the phenotypic differences in IHP1 versus ILP1 root architecture.

**Figure 1.**
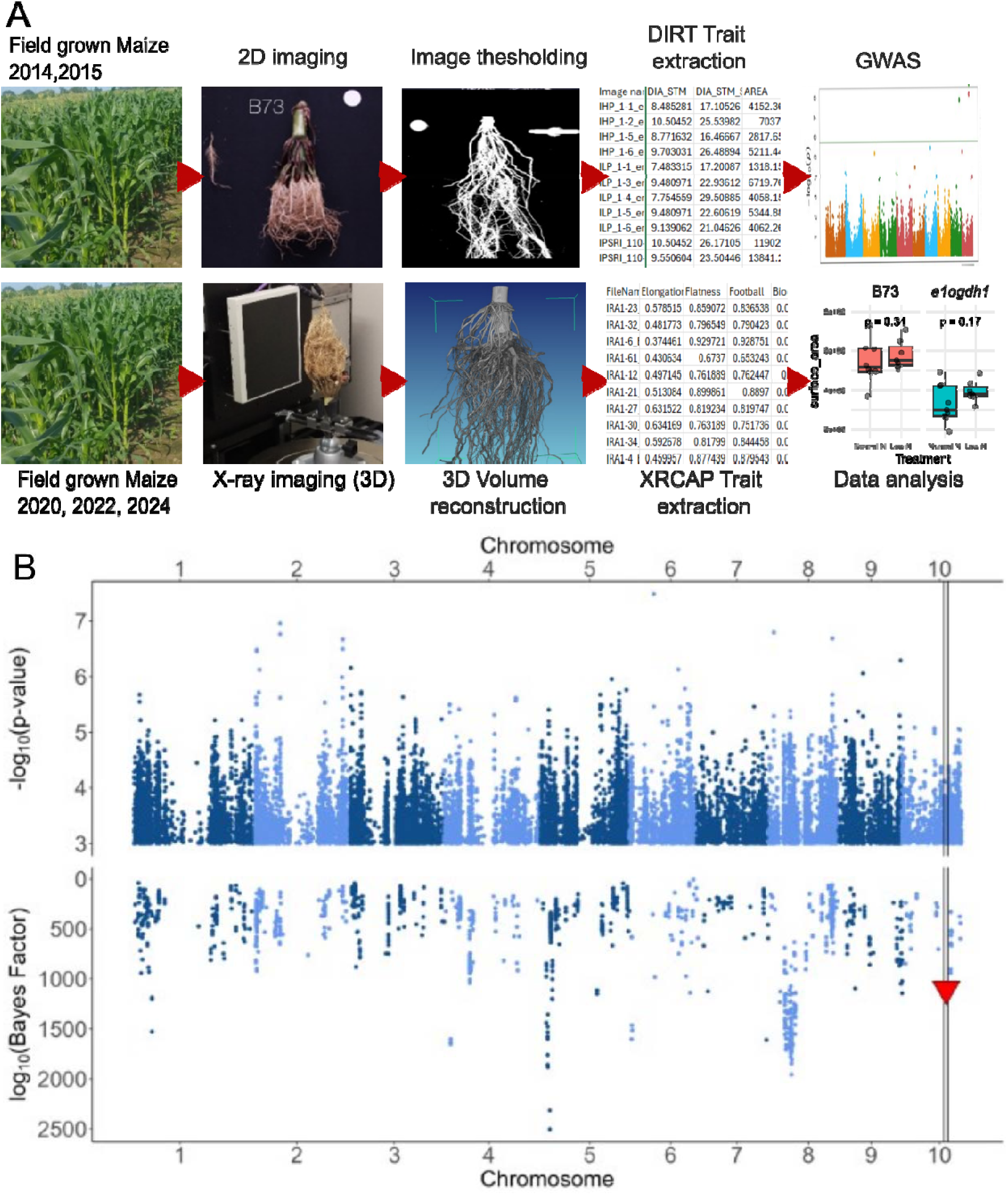
Several root trait-associated SNPs (TAS) were indirectly selected during the over 100 years of selection for grain protein (A) Data extraction and analysis workflow. (B) Manhattan plot (top) and Mashattan plot (bottom) of the 2014 and 2015 root crown GWAS using root architectural traits. The top SNP in the QTL is denoted as a red triangle.

### E1OGDH1 identified as a candidate gene in the QTL region

Out of the several loci identified by Multivariate Adaptive Shrinkage (MASH) analysis, we focused on a quantitative trait locus (QTL) on chromosome 10 (Fig. 2A). This region was consistently associated with three distinct root traits: stem diameter (DIAMETER), root distribution along the y-axis (RDISTR_Y), and skeletal depth (SKL_DEPTH, Fig. 2B) which had a heritability of 18.6%, 12.9%, and 11.8% respectively (Das et al. 2015). Notably, a significant SNP within this region, identified in the 2014 dataset, was located in the proximal upstream region of E1OGDH1, which encodes the E1 subunit of the 2-oxoglutarate dehydrogenase complex, a key enzyme mediating nitrogen assimilation and carbon balance. Subsequent MASH analysis integrating GWAS results from both 2014 and 2015 confirmed the presence of a QTL encompassing E1OGDH1 within a region of high linkage disequilibrium (Fig 2A). Given known differences of nitrogen assimilation and free amino acid accumulation between IHP and ILP (Hoener and DeTurk, 1938; Lohaus *et al*., 1998) we prioritized this candidate gene due to its potential to link nitrogen and protein metabolism pathways to root system architecture.

**Figure 2.**
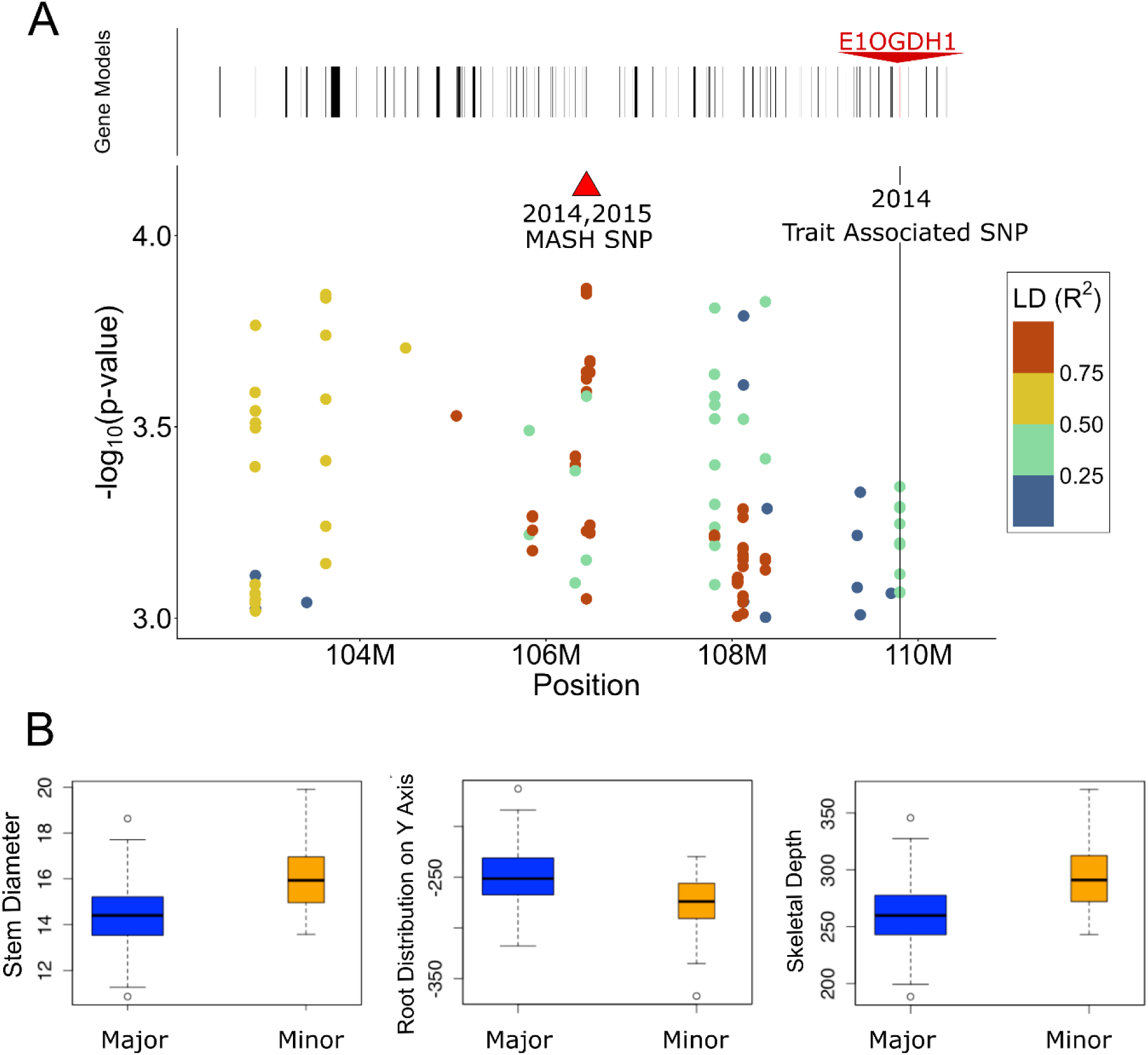
E1OGDH1 is a candidate gene involved in root system architecture and nitrogen relations. **(A) Local linkage disequilibrium (LD) structure surrounding the QTL region on chromosome 10.** The local region around the MASH significant peak, which is marked with the red triangle. SNPs within the LD region are represented as dots and colored based on their LD. The LD values are the R² to the MASH identified SNP. Physical locations of gene models in this region are plotted as black rectangles above the LD plot. The specific SNP linked to *E1OGDH1* is indicated with the vertical line. (B) Genetic effects of the major and minor allele for the three traits associated with the chromosome 10 QTL identified in 2014: Diameter of Stem (DIA_STM_SIMPLE, left plot), Root Distribution in the Y axis (RDISTR_Y, middle plot), Skeletal Depth (SKL_DEPTH, right plot). The width of the boxes is proportionate to the number of IPSRI lines with either the major or the minor allele. Trait definitions are provided in (Das *et al*., 2015).

E1OGDH1 occupies a key position at the intersection of carbon and nitrogen metabolism. Its substrate, 2-oxoglutarate, serves dual roles: as an intermediate in the tricarboxylic acid (TCA) cycle for energy production, and as a carbon skeleton for nitrogen assimilation via the glutamine synthetase/glutamate synthase (GS/GOGAT) pathway (Masclaux-Daubresse *et al*., 2006). Emerging evidence also suggests that glutamate, a product of this pathway, when applied exogenously has been shown to repress primary root elongation while enhancing lateral root development in Arabidopsis (Ma *et al*., 2022). Given the central role of E1OGDH1 in these metabolic and signaling networks and the lack of prior characterization regarding its influence on root system architecture, we selected this gene for further investigation.

### Promoter variation in E1OGDH1 defines IHP1 and ILP1 alleles and is co-incident with expression differences in root tissue

To investigate further whether E1OGDH1 could be the gene responsible for the identified QTL, we compared the sequence of the promoter and coding regions of E1OGDH1 in the IHP1 and ILP1 inbreds and several IPSRI lines from the core population. PCR amplification based on primers designed from B73 or other modern reference germplasm was repeatedly unsuccessful. Therefore, we turned to nanopore long-read technology to sequence the gene region. The IHP1 sequence was identical to B73 except for 6-bp indel 451-bp upstream of the transcription factor start site and 2 intronic SNPs. The only coding region difference between IHP1 and ILP1 was a single conservative amino acid substitution (leucine997 to methionine) in the last exon of ILP1 (Fig.3A). Structural modeling using mCSM <u>(Pires et al. 2020)</u> predicted only mild destabilization (ΔΔG = -0.615 kcal/mol) and negligible binding affinity change (ΔΔG = -0.49 kcal/mol). Structural localization confirmed that position 977 resides in the C-terminal E1 main region distant from both the ThPP-binding active site and the N-terminal DSBD motif responsible for E1-E2 subunit interactions, a result consistent with the poor evolutionary conservation at this position observed in BLAST alignment against the human homolog <u>(Zhang</u> <u>et al. 2024)</u>. Collectively these findings indicate that the amino acid difference is unlikely to affect E1OGDH1 enzymatic activity or multienzyme complex assembly. However, multiple polymorphisms were found in the surrounding sequence. In the 3’ region of ILP1, an appx. 4,213-bp transposon and 140-bp sequence inserted just downstream of the 3’UTR. The 5’ putative promoter region of ILP1 contained two indels within 600-bp of the transcription start site relative to the IHP1 and B73 alleles (Fig. 3A): an approximately 6660-bp transposon insertion 3260bp upstream and a 116-bp MITE transposon 365bp upstream. We took advantage of the MITE polymorphism to design genotyping primers that distinguished IHP1 from ILP1 E1OGDH1 alleles (Methods) and allowed us to track their segregation in the IPSRI population inbreds. Since the coding region of E1OGDH1 had no apparent functional difference between IHP1 and ILP1, we hypothesized that the promoter variation could be causing differences in E1OGDH1 expression that might be affecting root system architecture. Total RNA extracted from root tissue of IHP1 and ILP1 plants grown in paper towels for two weeks revealed that the ILP1 allele showed decreased expression of E1OGDH1 relative to the IHP1 allele in both nodal and lateral roots, with a nearly 3.5-fold decrease in laterals (Fig. 3B). This result is consistent with the transposon insertions weakening the promoter activity of the ILP1 E1OGDH1 allele, which could underlie the observed phenotypic effects, although we did not directly test this hypothesis in this study.

**Figure 3.**
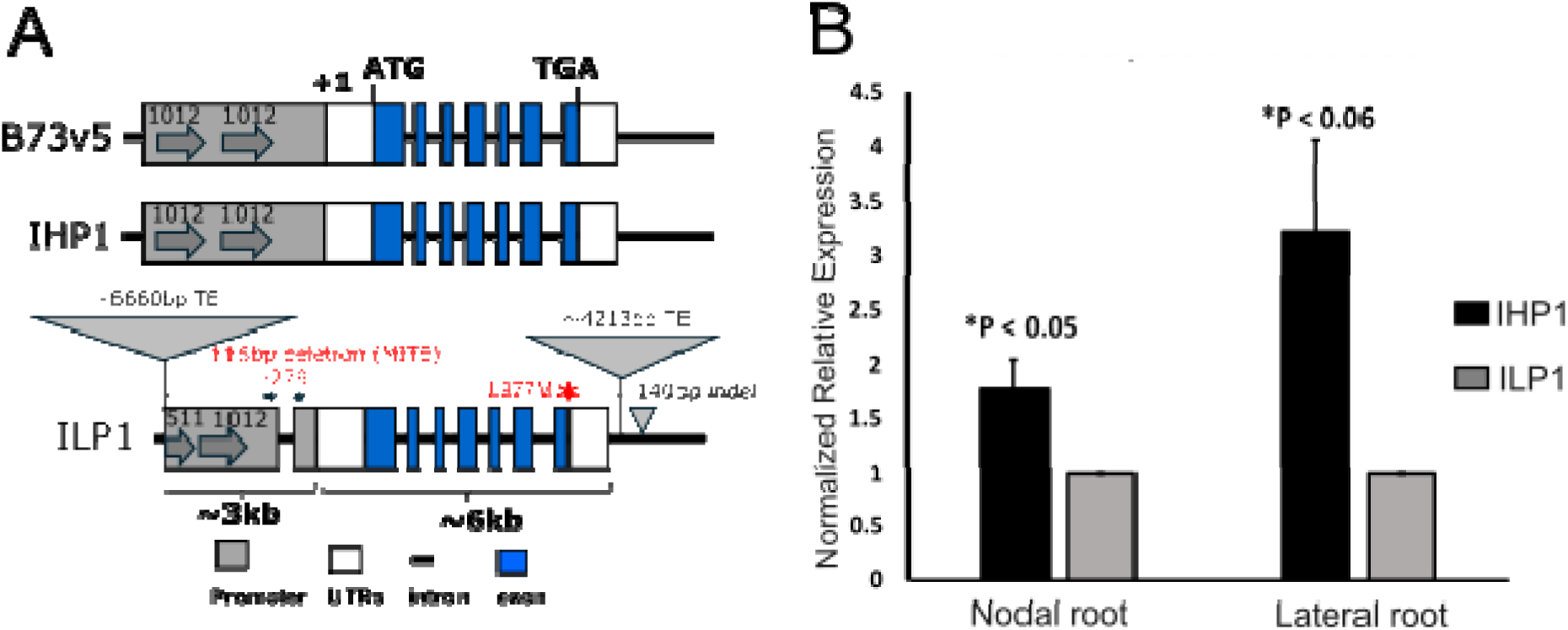
Sequence and RNA expression variation among different E1OGDH1 alleles. (A) Schematic of the promoter and coding regions of E1OGDH1 from B73, IHP1 and ILP1. The B73 and IHP1 alleles both harbor a 1012-bp direct repeat (arrow) and are identical except for a 6-bp indel 451bp upstream of the transcription start site. ILP1 harbors three TE insertions in flanking regions relative to IHP1 and B73, the MITE region used for genotyping is colored in red. The amino acid change is shown in the red asterisk. (B) E1OGDH1 expression level in IHP1 and ILP1 in nodal and lateral root tissue. Expression level is normalized to ILP1, which is set to 1.

### Promoter variation in E1OGDH1 is linked with root trait variation across two years of field experiments

We next sought to test whether allelic variation in the promoter region of E1OGDH1 is specifically associated with root system architecture differences. Unfortunately, we could not identify an existing mutant resource in maize that would allow us to directly make this determination in an otherwise homogeneous background. Instead, employed the genotyping assay developed for the promoter located 116-bp MITE indel and asked whether root system architecture traits were significantly different among IPSRI inbred lines homozygous for either the IHP1 or ILP1 E1OGDH1 allele. To maximize our chances of detecting subtle RSA phenotypic effects, we employed 3D X-ray tomography (XRT) imaging and analysis, which increases information content and is more sensitive to detection of subtle differences in root phenotypes (Bray and Topp, 2018; Shao *et al*., 2021; Hein *et al*., 2025).

We conducted field experiments in two years (a third was ruined by a catastrophic flooding event) to test this hypothesis. The 2020 experiment included 113 lines from the IPSRI population. Of these, 62 lines were homozygous for the IHP1 allele at the E1OGDH1 locus and 42 lines were homozygous for the ILP1 allele, with the remaining lines being heterozygous or otherwise unclassified at this locus. A total of 230 root crowns were phenotyped: 137 from plants carrying the IHP1 allele and 93 from plants carrying the ILP1 allele. Several traits exhibited significant differences (Supplementary Table 1), with solidity being particularly notable. Solidity is a measurement that estimates how thoroughly a root system explores a volume of soil, which is calculated by the soil the root crown encompasses divided by the biomass of the root crown. Lateral root branching density has been previously associated with root responses to low external nutrient conditions (Postma, Dathe and Lynch, 2014; York and Lynch, 2015), suggesting a potential link between metabolism, nitrogen sensing, and root system “foraging” for soil nitrogen resources. We observed that IPSRI lines containing the ILP1 allele had significantly higher solidity values than IPSRI lines containing the IHP1 allele (t-test p-value = 0.001, Fig. 4A).

**Figure 4.**
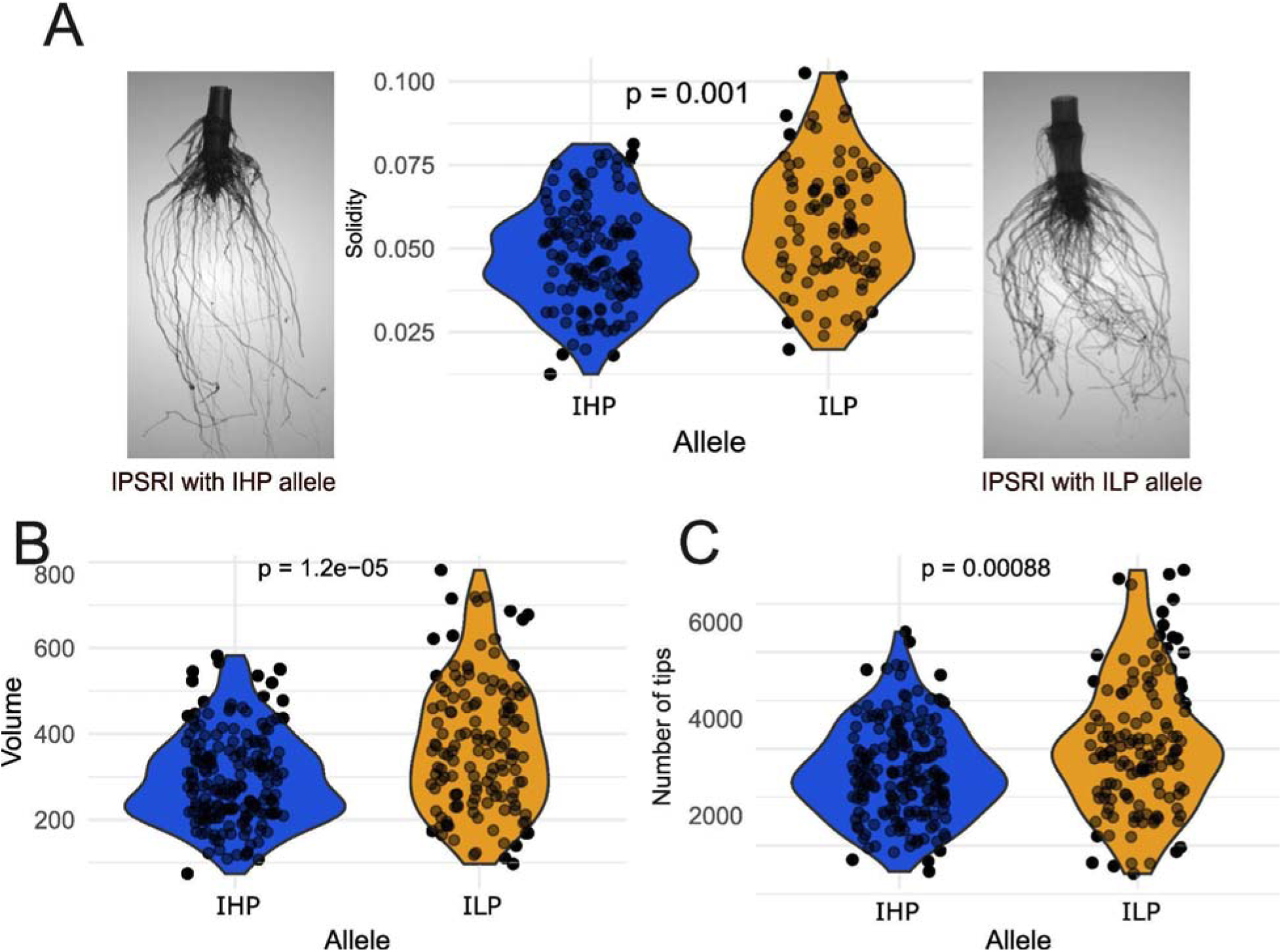
IHP and ILP alleles of E1OGDH1 correlate with different root traits in field studies. (A) Violin plot of root solidity for plants with IHP and ILP alleles of E1OGDH1 in 2020. 3D reconstructions of representative IPSRI root crowns are shown on the sides: left, IPSRI with the IHP allele; right, IPSRI with the ILP allele. (B) Violin plot of root volume for plants with IHP and ILP alleles of E1OGDH1 in 2022. (C) Violin plot of root tip number for plants with IHP and ILP alleles of E1OGDH1 in 2022. Wilcoxon test p value between allele group is shown on plots. Each dot represents a root crown sample.

In 2022, we repeated the experiment using a subset of 14 IPSRI lines that were genotyped as homozygous for either the IHP1 or the ILP1 E1OGDH allele from the 2020 experiment, analyzing a total of 401 root crowns, of which 213 carried the IHP1 allele and 188 carried the ILP1 allele. A Wilcoxon test, revealed significant allelic effects for several traits, including volume and root tip numbers (Fig. 4B, C; Supplemental Table 1). Lines with the ILP1 allele showed increased root crown volume, suggesting greater biomass accumulation, as well as a higher number of root tips, indicating increased root branching. Together, these results provide genetic evidence that E1OGDH1 variation can influence root system architecture and supports E1OGDH1 as a strong candidate gene underlying the chromosome 10 QTL we identified in our initial field root phenotyping campaign.

### An E1OGDH1 knockout mutant in a B73 inbred background shows whole plant phenotypes and an altered root growth response to external nitrogen

The results from the IPSRI population provided insight into the role of E1OGDH1 in root system architecture. However, explicit testing of allelic effects on plant growth and physiology is limited in this non-homogenous background derived from the historical heirloom variety Burr’s White. We therefore generated CRISPR mutants in the well studied B73 background. The guide RNA (sgRNA) of the CRISPR construct targeted the first exon of the E1OGDH1 and its paralog E1OGDH2. Several mutants were generated from this CRISPR construct, including a 1bp deletion, a 12bp deletion, and a 24bp deletion mutant in E1OGDH1 and a 1bp insertion in E1OGDH2 (Supplementary Fig. 2). We decided to focus on the 1bp *e1ogdh1* deletion mutant, which induced a frameshift mutation that truncated the predicted protein of 1025 amino acids at the amino acid 207 position and encoded additional 66 amino acids unrelated to the original gene product (Fig. 5A), almost certainly rendering the enzyme non-functional. Given that the gene is involved in an important crossroad between energy generation and nitrogen assimilation, we expected pleiotropic growth defects above- and belowground, but hypothesized that the mutant would have a different relative response to external nitrogen availability that could provide clues to its role in root system growth and function.

**Figure 5.**
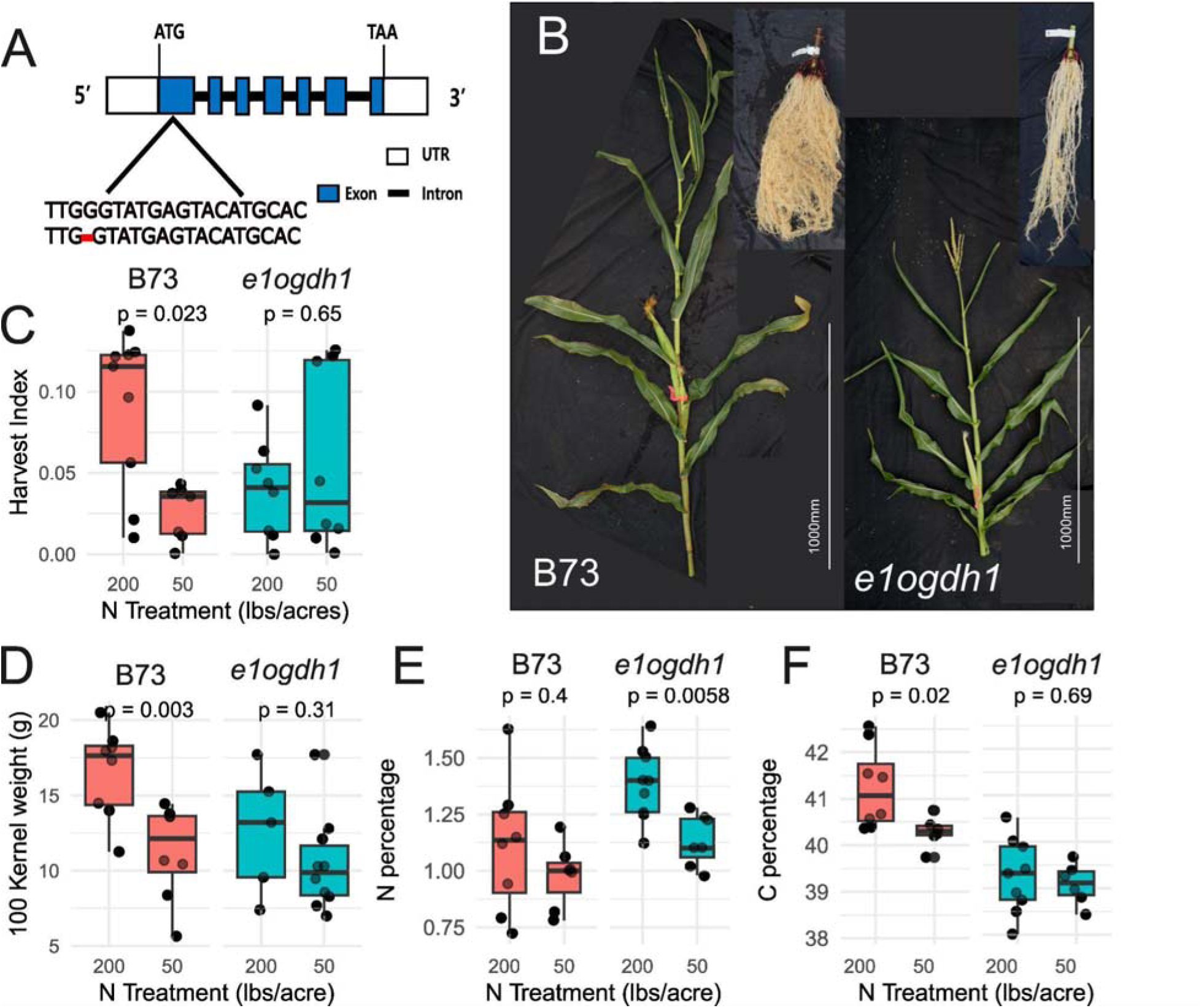
CRISPR mutant of *e1ogdh1* exhibits altered plant size and developmental phenotypes. (A) schematic of the 1bp mutation of E1OGDH1 generated from CRISPR-Cas9. (B) Shoot (left) and root (upper right) of mature B73 and *e1ogdh1.* Boxplot for B73 and *e1ogdh1* in the field under different nitrogen treatments. (C) Harvest index. (D) 100 kernel weight. (E) Shoot N percentage (%). (F) Shoot C percentage (%). Student t-test p-value between N treatment is shown in the plot. The sample size for each box is between 7 and 9. Each dot represents a root crown sample.

We conducted a regulated field trial with four nitrogen levels for wild-type B73 and the e1ogdh1 mutant. For clarity, we present results from only the highest (“standard” 200lbs/acre, 224kgs/hectare) and lowest (50 lbs/acre, 56kgs/hectare) nitrogen levels. At all levels, *e1ogdh1* plants exhibited reduced stature relative to B73, including reduced plant height and smaller root systems (Fig. 5B, not shown). As expected, the harvest index of B73 (percent grain weight relative to aboveground biomass) increased with greater soil nitrogen availability (Fig. 5C, Supplementary Table 2), whereas *e1ogdh1* showed little change across nitrogen levels. Similarly, grain production, as measured by 100 kernel weight, increased as expected in B73 with increased soil nitrogen but not in *e1ogdh1*. Analysis of shoot tissue composition revealed that under normal nitrogen conditions, *e1ogdh1* showed significantly higher relative nitrogen content than B73 (1.38% vs 1.11%, Welch’s t-test p = 0.042) and significantly lower carbon content (39.34% vs 41.24%, Welch’s t-test p = 0.0004) compared to B73, indicating an altered carbon-nitrogen balance in the mutant consistent with impaired 2-oxoglutarate dehydrogenase activity. Under low nitrogen, B73 maintained its N homeostasis while *e1ogdh1* significantly decreased tissue N, albeit at an elevated level to B73 (Fig 5E,F, Supplementary Table 2). Overall, *e1ogdh1* plants were smaller, produced fewer leaves, maintained a consistently low harvest index and grain production regardless of nitrogen supply, and failed to maintain a consistent nitrogen concentration within the plant under nitrogen stress. These findings collectively indicate that E1OGDH1 is required for normal physiological responses to soil nitrogen availability.

To evaluate the impact of *e1ogdh1* on root system architecture and growth responses to soil nitrogen availability, we evaluated soil excavated root crown phenotypes by 3D X-ray analysis (Fig. 6, Supp Table 3). The mutation had a strong influence on root volume, a proxy for biomass, with surprisingly little effect of soil nitrogen in either genotype (Fig. 6A). The number of root tips (an estimate of lateral root abundance) showed a reduction in *e1ogdh1* but both genotypes responded similarly to low N by increasing root system branching (Fig. 6B). The width-to-depth ratio describes the overall shape strategy of the root crown, indicating whether it predominately spreads laterally to forage in shallow, nutrient-rich surface soil or grows deeper to access water and mobile nutrients like nitrate at lower soil horizons. Horizontal maximum root width (HorEqDiameter) captures the maximum lateral spread of the root crown and similarly reflects how broadly the root system occupies horizontal soil space. As expected, both traits were influenced by soil nitrogen in B73 with a shift to wider, shallower root systems, but this response was dampened in *e1ogdh1* mutants with a statistically non-significant trend in the same direction as B73 (Fig. 6C,D). Average edge length describes the distance between lateral roots, akin to lateral root branch density. The *e1ogdh1* mutation had the largest effect on this trait, with less dense lateral rooting under standard N conditions, but a significant increase under low N, in contast to B73 which maintained a constant relative value (Fig. 6E). This may have contributed to the trend toward increasing solidity in low N for *e1ogdh*1 as additional lateral root would increase exploraiton thoroughness in without significant increases in root system width or depth (Fig. 6C,E).

**Figure 6.**
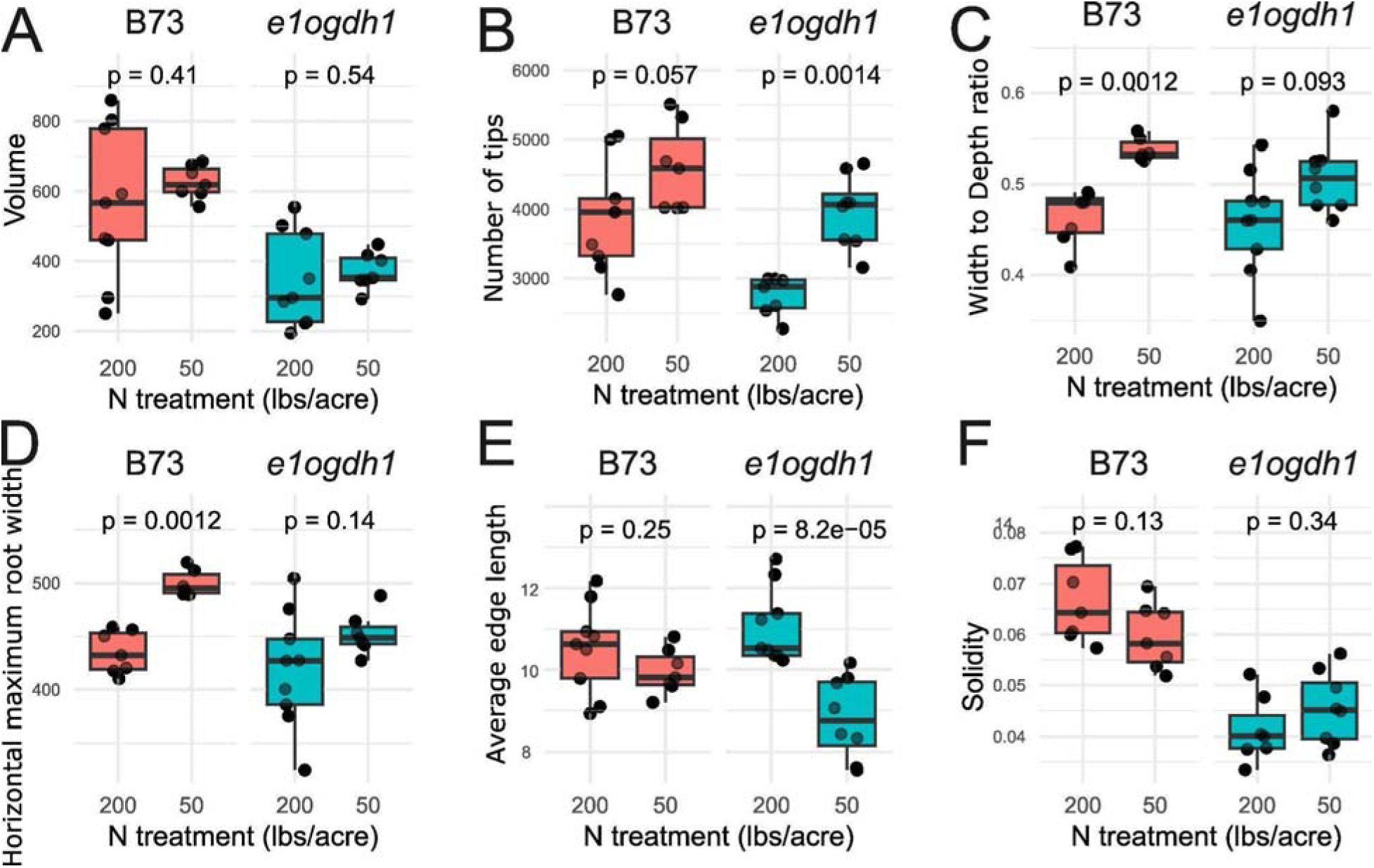
B73 and *e1ogdh1* root traits show significant differences by genotype, Nitrogen treatment, and genotype-nitrogen interaction. Boxplots of root traits of B73 and *e1ogdh1* in the field with different nitrogen treatments. (A) Volume. (B) Root tip numbers. (C) Width-to-depth ratio. (D) Maximum Diameter (HorEqDiameter). (E) Average edge length. (F) Solidity. Each sample n =7∼9. Statistical Student’s t-test significance between nitrogen treatments is shown on plots.

To further evaluate genotype-by-treatment interaction effects, we conducted ART ANOVA, (Supplementary Table 4). Among the traits showing strong interaction effects, the average edge length, which is the length between each branching point and estimates branching density, showed genotype-dependent responses to nitrogen treatment (Fig. 6E, genotype-by-treatment interaction ART ANOVA p-value 0.0054, p <0.01). B73 showed a 5.59% decrease under nitrogen stress conditions, while *e1ogdh1* had a substantial 25.32% decrease, demonstrating more branching in the mutant compared to the wild type. Similarly, solidity displayed a significant interaction effect (Fig. 6F, genotype-by-treatment interaction ART ANOVA p-value 0.0254, p<0.05). B73 had an 11.74% decrease in solidity under nitrogen stress, whereas *e1ogdh1* had a 9.25% increase.

These findings collectively indicate that E1OGDH1 contributes to root system architecture and growth responses to soil nitrogen availability. The influence is allometric - under nitrogen stress, B73 tends to develop wider overall root systems, suggesting a topsoil foraging strategy, while *e1ogdh1* increases lateral rooting (Figure 7). This distinction is further supported by the differences in traits such as average edge length and solidity across nitrogen treatments. These changes may in turn affect relative nitrogen uptake rates, as nitrogen tissue concentration was elevated in e1ogdh1 mutant plants despite their small size (Fig. 5E). Overall, these results support that E1OGDH1 plays a key role in mediating root architectural responses to nitrogen availability, potentially linking primary metabolism to environmental growth adaptation.

**Figure 7.**
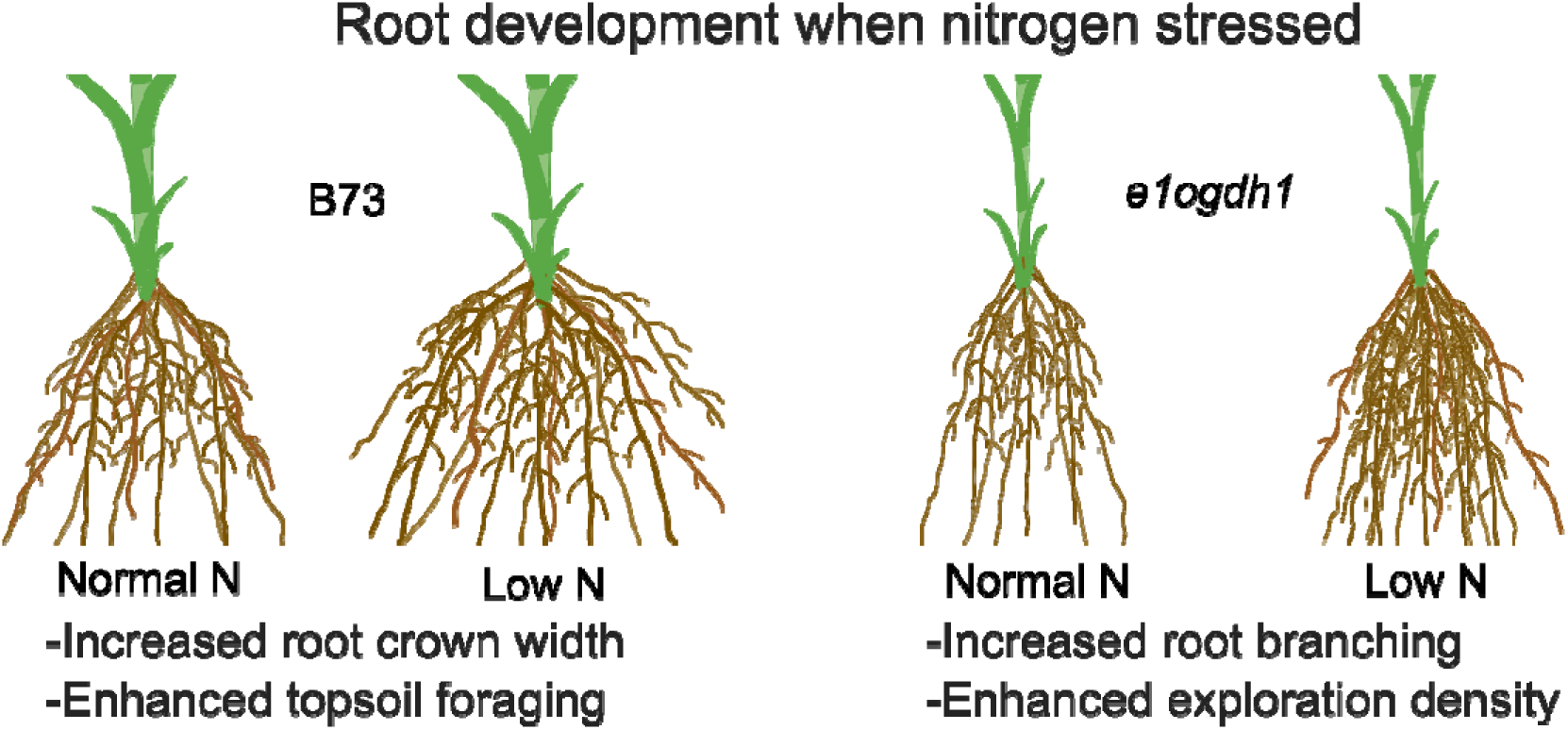
Model of B73 and *e1ogdh1* response to Nitrogen stress. To mitigate nitrogen stress, B73 mainly increases the width of the root to enhance topsoil foraging. In contrast, *e1ogdh1* employs a lower energy strategy that increases branching and enhances exploration of the occupied soil space.

### Loss of E1OGDH1 leads to changes in root architectural genes and nitrogen responsive genes in normal and nitrogen stress conditions

To further understand potential mechanisms linking disruption of a carbon-nitrogen metabolism gene to changes in root system architecture and soil nitrogen relations, we conducted an RNAseq experiment in roots under both normal and nitrogen-limiting conditions. Aboveground (aerial) nodal root tips that reached the soil and actively growing were collected from R1 stage *e1ogdh1* and B73 plants grown under high and low nitrogen conditions, and RNA was extracted for transcriptomic analysis. We focused our analysis on a curated list of 205 genes of interest, comprising genes associated with root phenotypes in MaizeGDB, annotated root-related genes from Ren *et al*., 2022, and nitrogen-related genes including nitrate and ammonium transporters and TCA cycle genes in the vicinity of E1OGDH1.

We first examined genes that showed significant differential expression between B73 and *e1ogdh1* under normal nitrogen conditions (Fig. 8, left panel). Of the 205 genes surveyed, 41 showed significant differential expression at high nitrogen. Several genes known to promote root biomass, including nrg7, NPF6.4, brd1, and ZmEXPA5, showed higher expression in B73 relative to *e1ogdh1*, while bhlh204 showed the opposite pattern with higher expression in the mutant (Ren *et al*., 2022; Zhang *et al*., 2022; Zhang, Chen and Yu, 2023; Li *et al*., 2024; Nedelyaeva *et al*., 2024; Qin *et al*., 2025). HKT1, a gene known to negatively affect stem diameter, was elevated in e1ogdh1, consistent with the reduced stem diameter observed in this genotype (Li *et al*., 2019). Genes regulating lateral root development (Z. Li *et al*., 2018; Li *et al*., 2025), including dlr2 and PIN1a, were also elevated in *e1ogdh1*. Several genes with reported roles in nitrogen metabolism, including pco133519, nnr4, AMT1.3, clc1, and LOC100279218, showed dampened expression in the mutant (Dechorgnat *et al*., 2019; Ma *et al*., 2020; Zhang *et al*., 2026). Among the nitrate transporters, ZmNPF4.2 and ZmNRT3.1B, which are normally expressed at low levels in roots, showed higher expression in *e1ogdh1* (Dechorgnat *et al*., 2019). In contrast, other nitrate transporters including NPF5.13, NPF8.8, NPF6.4, NPF8.10, NPF1.3, NPF5.2, NPF8.15, NPF5.15, and ZmNRT2.5 showed significant expression differences between B73 and *e1ogdh1* (Jia *et al*., 2023), differing in both direction and magnitude, suggesting that nitrogen transporter regulation is altered in *e1ogdh1* even under nitrogen-sufficient conditions. Among genes encoding other TCA cycle enzymes, idh3 and idh5 for isocitrate dehydrogenase that supplies substrate for E1OGDH1 showed elevated expression in *e1ogdh1* at high nitrogen, which may reflect a response to altered TCA flux.

**Figure 8.**
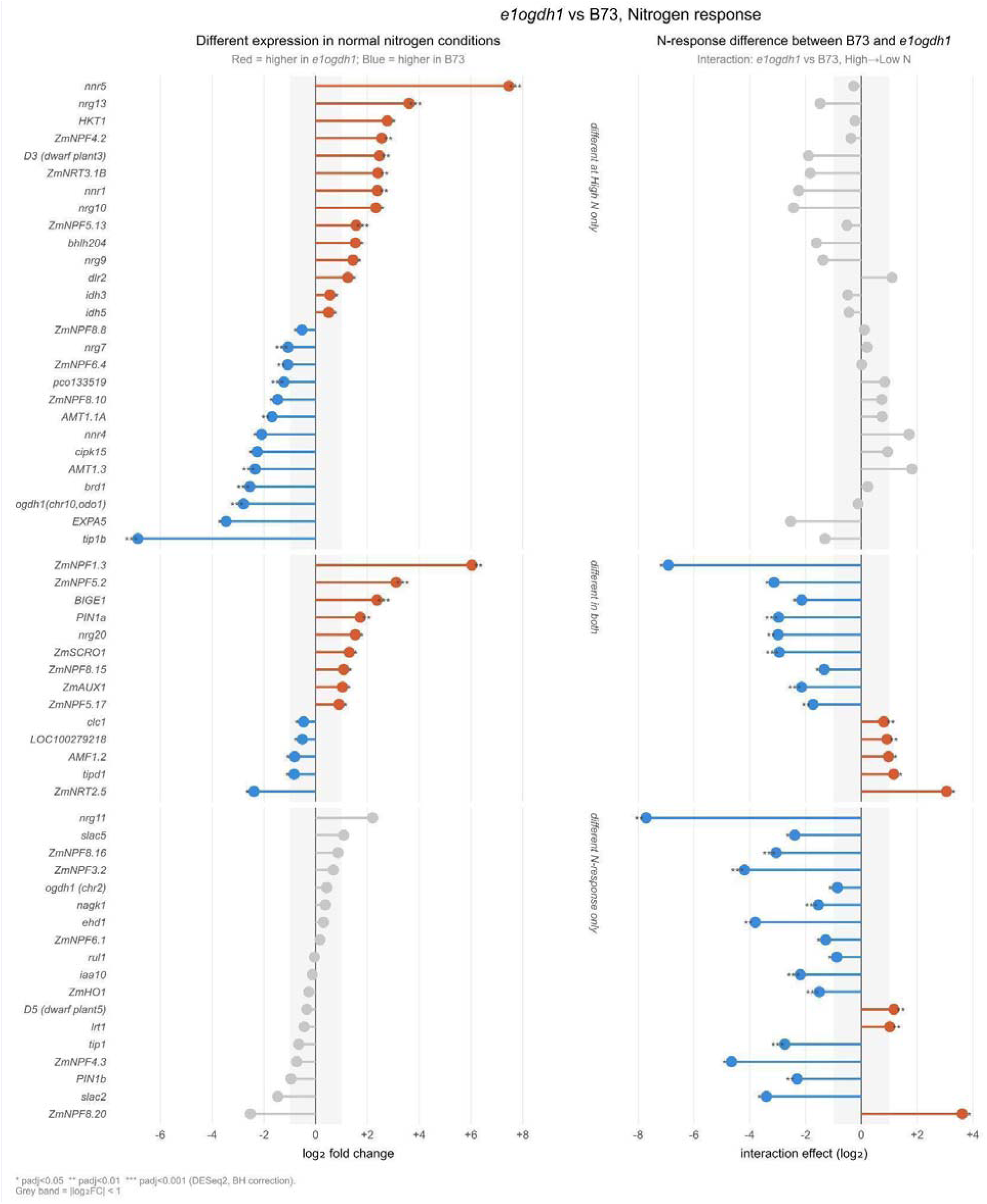
Root- and nitrogen-related genes show differential expression. (Left) Baseline expression differences between *e1ogdh1* and B73 under high nitrogen. Each point represents one gene, with bar length indicating the magnitude of the genotype effect (log□ fold change of *e1ogdh1* relative to B73) at high nitrogen in root tissue. Red points indicate higher expression in *e1ogdh1*; blue points indicate lower expression in *e1ogdh1*. The grey shaded band indicates |log□FC| < 1. Genes are grouped by expression pattern: group A, significant difference at high nitrogen only; group B, significant difference in nitrogen response only; group C, significant difference in both contrasts. Significance was assessed using DESeq2 Wald test with Benjamini-Hochberg correction; asterisks indicate padj < 0.05 (*), padj < 0.01 (**), and padj < 0.001 (***). (Right; interaction effect) Differential nitrogen response between *e1ogdh1* and B73 in roots. Bar length represents the difference in nitrogen response between genotypes, calculated as log□FC(lowN/highN) in *e1ogdh1* minus log FC(lowN/highN) in B73. Red points indicate a greater induction or attenuated repression under low nitrogen in *e1ogdh1* relative to B73; blue points indicate a stronger repression or attenuated induction in *e1ogdh1* relative to B73. Gene grouping and significance thresholds are as in panel A. The grey shaded band indicates |interaction log□FC| < 1. Gene names are shown on the y-axis; genes are ordered within each group by baseline fold change.

We then examined genes showing differential nitrogen responses between mutant and WT (Fig. 8, right panel). Thirty-two genes showed a significantly different transcriptional response to nitrogen limitation in *e1ogdh1* compared to B73, with the majority exhibiting a dampened response to nitrogen stress in the mutant. Among genes showing differences in both baseline expression and nitrogen response, the predominant pattern was opposite directional responses between genotypes: B73 induced these genes under low nitrogen while *e1ogdh1* repressed them, or vice versa. This was particularly evident for the auxin transport genes PIN1a and ZmAUX1 and the nitrate transporter ZmNPF5.17, suggesting that *e1ogdh1* disrupts both the constitutive and nitrogen-regulated expression of genes involved in root architecture and nutrient acquisition. BIGE1, a gene known to negatively affect lateral root number, showed higher expression in *e1ogdh1* under normal nitrogen conditions but was strongly induced in B73 under nitrogen stress, consistent with the lateral root phenotype observed in this study (Suzuki *et al*., 2015). Several other genes with established lateral root phenotypes, including ZmHO1, lrt1, tip1, and PIN1b, also showed altered nitrogen responses in *e1ogdh1* (Han *et al*., 2012; Z. Li *et al*., 2018; Zhang *et al*., 2020; Baer *et al*., 2023). Among genes that did not differ significantly under normal nitrogen conditions but diverged under nitrogen stress were genes associated with greater root biomass, such as, rul1, and iaa10 (Ren *et al*., 2022; Singh *et al*., 2024). These genes showed significantly stronger induction in B73 than in *e1ogdh1* under low nitrogen. Similarly, slac2 and slac5, genes associated with root pulling force, showed attenuated nitrogen responses in *e1ogdh1* relative to B73 (Feng *et al*., 2025). Strikingly, genes including PIN1b, iaa10, ehd1, and ZmHO1 showed strong nitrogen-responsive induction in B73 that was completely absent in *e1ogdh1*, indicating that the E1OGDH1 mutation can specifically impair the transcriptional response to nitrogen limitation for a subset of genes without affecting their basal expression levels. The enrichment of auxin-related genes in this category, including PIN1b and iaa10, suggests that the *e1ogdh1* mutation may disrupt the coordination between nitrogen signaling and auxin-mediated root development. Furthermore, several genes showed opposite directional responses between genotypes under low nitrogen, induced in B73 but repressed in *e1ogdh1*, including ZmNPF4.3, slac2, and tip1, pointing to fundamental differences in how the two genotypes reprogram gene expression in response to nitrogen limitation. Taken together, the altered expression of genes regulating root size, lateral root development, auxin transport, and nitrogen acquisition in *e1ogdh1* provides a transcriptional basis for the root phenotype observed in this mutant and suggests that E1OGDH1 is required for the normal coordination of carbon-nitrogen metabolism and nitrogen-responsive root development in maize.

## Discussion

Improving nitrogen use efficiency in maize requires a deeper understanding of how plants coordinate internal nitrogen metabolism with developmental responses in the root system. Root system architecture (RSA) determines where and when plants encounter soil nitrogen, yet the genetic and metabolic mechanisms linking internal nitrogen status to dynamic root growth remain largely unresolved in maize and other crops (Forde, 2002; Gallais and Hirel, 2004; Krapp *et al*., 2011). Here, we combine multi-year field root phenotyping, quantitative genetics, and genome editing to identify E1OGDH1 as a previously uncharacterized quantitative genetic regulator of maize RSA and nitrogen-responsive root development. Our results place this gene at the nexus of energy metabolism, nitrogen assimilation, and adaptive root architectural plasticity.

Our GWAS results demonstrate that image-derived root traits can be effectively used to identify new genes involved in RSA (Fig. 1C). Among the QTL identified, 66.78% were only identified in a single year, highlighting the strong influence of environmental effects on root development. This emphasizes the importance of conducting multi-year field experiments and integrating results across years using methods such as MASH, which can help identify genes like E1OGDH1 that were consistently associated with RSA in multiple years. Out of the multiple QTLs we identified, we focused on one in chromosome 10 that contained E1OGDH1 in its LD region. Although E1OGDH1 had not been identified previously as a root system architecture gene, it is part of the OGDH protein complex, which is known to be a rate-limiting step of the TCA cycle. This step serves as a critical metabolic branching point, determining whether the substrate 2-oxoglutarate proceeds through the TCA cycle or is redirected for nitrate assimilation and amino acid metabolism. Both pathways also supply intermediates to formation of other metabolites known to alter growth and nutrient responses in roots. Thus, E1OGDH1 became a compelling candidate for understanding the coordination between metabolic signaling and root development.

We then showed that both natural and artificial alleles of E1OGDH1 correlate with variations in root system architecture. Promoter transposon insertions are co-incident with reduced E1OGDH1 expression in ILP1 compared to IHP1 (Fig. 3) and these promoter variants were associated with variation in multiple root traits for the IPSRI population grown in the field across two years (Fig. 4). However, we did not detect strong correlations between grain protein concentration and root phenotypes in the IPSRIs, and E1OGDH1 alleles show no significant association with seed composition. Although some studies have shown that selection for grain traits can shape belowground phenotypes (York and Lynch, 2015; Messina *et al*., 2021; Ren *et al*., 2022; Sciarresi *et al*., 2025), here the role for E1OGDH1 in shaping RSA appears to be indirectly related to seed protein.

To further investigate E1OGDH1 function in a more modern and better characterized genetic background, we generated CRISPR mutants of E1OGDH1 in the B73 inbred. While we unfortunately did not recover knockdowns that mimicked the ILP allele, or other partially functional alleles, CRISPR knockout mutants demonstrated that E1OGDH1 is required for normal vegetative growth and nitrogen responsiveness. As expected for disruption of a central metabolic enzyme, mutants showed reduced shoot biomass and stature (Supplementary Fig. 2). Importantly however, loss of E1OGDH1 altered root architectural responses to nitrogen availability. Under nitrogen limitation, wild-type B73 increased root system width, consistent with an extensive foraging strategy that supports nitrate uptake. In contrast, *e1ogdh1* mutants increased branching and root system density, favoring exploitation of occupied soil volume over horizontal expansion (Fig. 6). This divergence indicates that E1OGDH1 helps determine the choice between exploratory and exploitative root growth strategies depending on energy status and nitrogen availability (Figure 8). The enhanced branching response of mutants echoes patterns observed in IPSRI lines with the low protein allele (ILP), suggesting a continuum between natural variation and full loss of function (Figure 4).

We then looked into the role of E1OGDH1 in root development and nitrogen response in maize by analyzing the transcriptional profiles of 205 curated genes in *e1ogdh1* and B73 roots under high and low nitrogen conditions. Our results reveal that disruption of E1OGDH1 broadly alters the expression of genes involved in root system architecture, nitrogen transport, nitrogen assimilation, and auxin signaling. An interesting pattern suggesting an attenuation of nitrogen-responsive gene regulation in the mutant implies that E1OGDH1, as a component of the OGDH complex, plays an important role in integrating TCA cycle flux with primary nitrogen assimilation and eventual transcriptional reprogramming in response to nitrogen availability. The observation that 41 genes showed significant expression differences between genotypes under normal nitrogen conditions suggests that loss of E1OGDH1 has broad consequences for root gene regulation even in the absence of nitrogen stress, which is observed in the root phenotypes. The idh3 and idh5 genes acting at the preceding step in the TCA cycle have previously been observed to be upregulated by broader abiotic stress (Wei et al., 2023), their elevated expression of in *e1ogdh1* under high nitrogen suggests the mutant may be experiencing metabolic stress due to energy depletion. Consistent with the root phenotype of *e1ogdh1*, genes promoting root biomass such as nrg7, brd1 and ZmEXPA5 showed reduced expression in the mutant, while genes associated with lateral root development such as dlr2 and PIN1a were elevated. The heterogeneous regulation of multiple nitrate transporters in opposite directions in *e1ogdh1* under high nitrogen suggests that loss of E1OGDH1 disrupts the coordinated regulation of nitrogen uptake rather than simply up- or downregulating nitrogen acquisition capacity.

E1OGDH1 is the primary route for synthesis of 2-oxoglutarate, the branch point between the TCA cycle and the GS-GOGAT pathway. Consequently, E1OGDH1 activity regulates the balance between respiratory energy metabolism and the supply of carbon skeletons for ammonium assimilation. The root phenotypes we observed are consistent with a model in which high E1OGDH1 activity increases flux through the TCA cycle, maintaining carbon energy balance and supporting thicker, less branched root systems suited to nitrogen-rich environments, whereas reduced E1OGDH1 activity elevates 2-oxoglutarate availability, promoting lateral root development and higher root density under nitrogen limitation. A key intermediary could be glutamate, which is considered to be a signaling molecule for general abiotic stress (Kan *et al*., 2017; Cheng *et al*., 2018; Zheng *et al*., 2018; Li, Ye and Qiu, 2019; Philippe *et al*., 2019). In Arabidopsis, the external application of glutamate to the root repressed primary root growth while promoting lateral root elongation (Walch-Liu *et al*., 2006; Forde and Walch-Liu, 2009), which stopped when returned to nitrate-rich conditions. These observations support a model in which glutamate and related metabolites may act as local cues to modulate root architecture in response to nitrogen availability, tuned by E1OGDH1 activity. Glutamate receptor-like channels (GLRs) unlike their mammalian counterparts have a wider range of substrates such as other amino acids, which could hint at their function in amino acid signaling in the roots (Qi, Stephens and Spalding, 2006; Vincill, Bieck and Spalding, 2012). However, some GLRs within the family still maintained a specific function for glutamate (Walch-Liu *et al*., 2006). GLR proteins responsive to extracellular glutamate could mediate this signaling, suggesting a molecular framework for root adaptation to metabolic and environmental cues (Forde *et al*., 2013).

The identification of *E1OGDH1* as a candidate gene influencing root system architecture and nitrogen metabolism is supported by recent evidence from Huang et al. (2026), who identified *THP3*, encoding glutamate-oxaloacetate transaminase 1 (GOT1), a key enzyme linking the TCA cycle to amino acid synthesis through reversible transamination between L-glutamate and oxaloacetate. Critically, GOT1 and the 2-oxoglutarate dehydrogenase complex encoded by *E1OGDH1* compete for the same α-ketoglutarate pool in root tissue, placing both enzymes at the same metabolic node connecting carbon and nitrogen metabolism. The functional importance of this crossroad is illustrated by our *e1ogdh1* knockout, which produced smaller plants and root systems alongside elevated whole-plant nitrogen percentage, a phenotype consistent with uncoupling of nitrogen assimilation from carbon metabolism and growth, and mirroring in the opposite direction the enhanced nitrogen assimilation, root biomass, and GS/GOGAT activities observed in the high-protein THP3 near-isogenic lines of Huang et al. (2026)

Although complete loss of E1OGDH1 compromises plant vigor, the discovery of natural promoter variation offers a promising avenue for allele tuning rather than full disruption. Crop improvement may benefit from modestly altering regulation or conditionally responsive alleles that enhance nitrogen capture under low-input systems while preserving biomass potential in fertilized conditions. Future work should explore evaluation of allelic series and tissue-specific regulation of E1OGDH1 and the introgression of potentially favorable haplotypes in elite germplasm if they don’t already exist. Additionally, more research on the impacts of E1OGDH1 regulation on root metabolites and expression of genes involved in nitrogen uptake and assimilation is warranted to better understand mechanisms linking soil nitrogen dynamics with internal nitrogen sensing and feedback to root system architecture. By identifying a core metabolic enzyme as a regulator of RSA and nitrogen sensing, this study underscores the potential utility of metabolically informed breeding strategies that complement traditional selection on yield and canopy traits. As global agriculture seeks to simultaneously increase productivity and reduce nitrogen inputs, genes like E1OGDH1 may offer promising targets for developing maize hybrids that use nitrogen more efficiently across variable environments and management regimes.

## Methods

### Plant Materials

The IHP1 and ILP1 inbred lines and the IPSRI population are from the Maize Breeding and Genetics Laboratory at the University of Illinois Urbana-Champaign and were used for the 2014, 2015, 2020, and 2022 field experiments. Genetic diversity of the IPSRIs was assessed using SNP marker genotypes from Dudley et al. (2007), with individual genotypes clustered using neighbor-joining implemented in TASSEL 5.0, and the dendogram visualized in using the webtool ITOL (Interactive Tree of Life, https://itol.embl.de/). Maize genotype B73 (PI 550473) and its edited lines were used in the 2024 mutant experiments.

### CRISPR-Cas9 gene editing for E1OGDH1 in maize B73

A CRISPR-Cas9 guide RNA (sgRNA) was designed and constructed with a pair of complementary oligos corresponding to the genomic sequence (5’-TTGGTATGAGTACATGCAC-3’) in the second exon of E1OGDH1 in an intermediate vector pENTR-gRNA1 before mobilizing the sgRNA cassette into a Cas9 expression vector through Gateway recombination following the protocol by (Char *et al*., 2017). B73 transformation with the CRISPR construct was implemented using the biolistic particle bombardment method along with a construct carrying the morphogenic regulator genes Baby Boom and Wuschel 2 (BBM/WUS2). The BBM/WUS2 construct was made based on the information as reported (Lowe et al. 2016). Transgenic plants were sequenced around the sgRNA region for the presence of edits and grown to maturity in a greenhouse. Transgenic plants were PCRed for the BBM gene and Cas9 gene to determine the present of the construct and the progeny plants to cross out the construct. Plants were sequenced for the CRISPR-Cas9 edits and to ensure homozygosity of desired edits.

### Field Experiments

Field experiments were conducted at multiple locations to evaluate the effects of E1OGDH1 allelic variation and nitrogen availability on root system architecture. In 2014 and 2015, the IPSRIs and other diverse maize genotypes were grown at the Crop Sciences Research and Education Center in Urbana, Illinois (latitude 40.08455, longitude -88.22483), on the University Illinois at Urbana-Champaign campus. The IPSRIs, IHP1, and ILP1 were grown in 2020 and 2022 at Plant Haven Research Farm in O’Fallon, Missouri (latitude 38.8487, longitude -90.6871). The B73 and e1ogdh1 mutations were grown in 2024 at Donald Danforth Plant Science Center Field Research Site in St. Charles, Missouri (latitude 38.8469, longitude - 90.4618).

#### 2014,2015 Field Experiment

The IPSRI inbreds were grown in field plots essentially as described in Cheng et al. 2021. Briefly, 20 seeds of each inbred were sown (May 8, 2014, or May 7, 2015) in a single row 5.3 m long with 76 cm row spacing. Genotypes were arranged by relative maturity to minimize variable shading effects. Nitrogen fertilizer was applied as granular ammonium sulfate banded at the root zone of V3-stage seedlings at a rate of 90 kg N per hectare. At physiological maturity after sampling for above-ground phenotypes, five root crowns were sampled from plants in the middle of each row. Grain protein concentration was measured from bulked seed of five ears from each plot using a Perten DA7200 near-infrared analyzer, using a custom calibration built to estimate the broad range of grain composition in the Illinois Long Term Selection Experiment.

#### 2020 Field Experiment

The 2020 experiment was conducted from June 5 to August 19, 2020, at O’Fallon, Missouri. 152 rows were planted using jab-type hand planters, including 140 IPSRI lines with three biological replicates each, and six rows each of IHP1 and ILP1 parental inbreds per replicate. Rows were 3.65 meters in length, with 1-meter row spacing and approximately 25.4 cm spacing within rows. The experiment was laid out in a randomized complete block design with 38 rows per range and 4 ranges per repetition. Ammonium nitrate was applied as the nitrogen source. At maturity, 230 root crowns were excavated, washed, and phenotyped using X-ray imaging and a custom analysis pipeline.

#### 2022 Field Experiment

The 2022 experiment took place at the same site in O’Fallon, Missouri, from June 14 to August 22, 2022. Pre-plant soil nitrogen levels were assessed via soil analysis. Two nitrogen treatments were implemented: high nitrogen (150 lbs/acre) and low nitrogen (50 lbs/acre), manually applied using urea. The experiment was laid out in a randomized complete block design of 14 lines of IPSRI with the IHP1 allele and 14 lines of IPSRI with the ILP1 allele. Spatial layout and root imaging protocols were consistent with the 2020 experiment.

#### 2024 Field Experiment

The 2024 field experiment was conducted at St. Charles, Missouri between May 31, 2024, and September 6th, 2024. Seeds were planted using a seed planter. B73 and *e1ogdh1* were planted 2 rows for each within each block, surrounded and flanked by B73 rows on each side. Each row is 4.57 m and 76.2cm within rows. 4 separate nitrogen levels, 50lbs/acre, 100lbs/acre, 150lbs/acre, 200lbs/acre, were treated by liquid nitrogen using a 4460 spray coup. Root imaging protocols were consistent with the 2020 experiment.

### Sample Collection

#### Root Crown Sampling for Image Analysis

At the silking stage R3, root crowns were excavated by digging approximately 30 cm around the base of individual maize plants and 30 cm deep using shovels. The collected roots were soaked in water and detergent solution, then thoroughly washed using high-pressure hoses within two days of excavation. Cleaned root crowns were subsequently dried for one week in a greenhouse before imaging.

#### Root Imaging and Image Processing

Dried root crowns were imaged using the North Star Imaging X5000 X-ray system. Two-dimensional X-ray projections were reconstructed into 3D volumes using efX-DR software (NSI, Rogers, MN). Following quality control checks of reconstructed volumes, root system traits were quantified using an internal 3D root phenotyping pipeline developed in our laboratory. Detailed descriptions of the image processing workflow and extracted root traits can be found in (Shao et al., 2021).

### Genotyping of plants

In brief, three 1/4 inch diameter leaf punches were collected from the youngest fully expanded leaf of the maize seedling 2 weeks after planting. DNA was extracted using a CTAB/Phenol/Chloroform method followed by ethanol precipitation. DNA was resuspended in 50uL H_2_O and quantified using a Nanodrop 2000 and diluted to 25-50 ng/μL. PCR reactions were set up using the GoTaq Green 2x Master Mix (Promega) according to the manufacturer’s instructions. Primers for genotyping are E1OGDH1_genotyping-F (CGAAAACAGGCAGAATTAGGTTAAA) and E1OGDH1_genotyping-R (CTGTGGGCAGCGCTTGTTTC). The primers target the 116bp indel in the promoter region of E1OGDH1. conditions were as follows: an initial 3 minute hold at 95°C, cycling conditions consisting of a denaturation step of 95°C for 30 seconds, an annealing step of 55°C for 30 seconds, and an extension step of 72°C for 45 seconds, repeated 34 times; and final extension at 72°C for 10 minutes followed by a 4°C hold.

### Nanopore sequencing

Leaf tissue was collected from 14-day-old IHP1 and ILP1 seedlings, frozen, and ground in liquid nitrogen, respectively. Long strands of high molecular weight DNA were extracted from ∼2 grams of the ground tissue in 10 mL of extraction buffer using the protocol in Clarke, (2009) with the following modifications: the buffer contained 2% PVP and 0.2% beta-mercaptoethanol. DNA was quantified using Qubit BR DNA reagent (ThermoFisher Scientific) and a DS-11 Fluorometer (DeNovix Inc.). Short DNA and RNA were removed with a Short Read Eliminator kit (Circulomics Inc.) following the manufacturer’s instructions. DNA was sheared with 20 passes through a 26G needle to obtain DNA fragments in the 10-50kb size range. DNA libraries were prepared using the Nanopore SQK-LSK109 Genomic DNA ligation kit and protocol. Libraries were sequenced on a Nanopore MinION sequencer using a FLO-MIN106 flow cell. The flow cell was treated with DNase for 30 minutes every 12-24 hours, when the number of available pores had dropped twofold, followed by a flush and reloading of the prepared library (Tyson, 2019). This enabled between 12-19 GB total read length, which provides ∼6-10x coverage of the maize genome. Basecalling was performed on fast5 output files from MinKNOW software using Guppy Basecaller v2.3.1 to create fastq files (Oxford Nanopore Technologies). Combined fastq files were mapped to B73 AGPv4 using minimap2 (Li, 2018). Samtools (Li *et al*., 2009) was used to convert SAM to BAM files, to sort BAM files, and to create a BAM index file. IGV software version 2.7.2 (Broad Institute) was used to view aligned reads and identify sequence variation between mutant and wild type. Indels and clipped reads in the promoter regions were used for BLAST on MaizeGDB’s ‘Maize Transposable Element Database – Wessler’ as well as RepBase to identify transposon insertions.

### RNA extraction and cDNA generation

Samples of emerged nodal roots and lateral roots along the primary root axis were harvested from IHP1 and ILP1 seedlings grown in controlled growth chamber conditions for 14 days after sowing the seeds in wet paper towels. Plants were grown under a 14-hour day length, 28°C day and 22°C night temperature, and 60% relative humidity. Nodal and visible lateral root tissues were collected separately from the same plant 14 days after moving to the growth chamber. Tissues for RNA extraction were collected and immediately frozen in liquid nitrogen. Tissue was ground while frozen in 2 mL Eppendorf SnapCap tubes with three 3.175 mm stainless steel balls in a Retsch MM400 until the tissue turned to powder. RNA extraction was performed with Trizol and a Zymo Direct-zolTM RNA Miniprep kit following the manufacturer’s protocol. cDNA synthesis was performed with the Takara PrimeScriptTM RT Reagent Kit. RNA and cDNA were quantified using a DeNovix DS-11 Fluorometer.

### RT-qPCR

Primer pairs were designed using primer3 with at least one primer flanking an exon-exon junction (Table S1). Reference genes were chosen from the list of genes in Manoli et al. 2012 or Lin et al. 2014 after comparing root tissue-specific expression from (Stelpflug et al., 2016). PCR reactions were made as follows in 10µL total volumes: 5 µL 2XSYBR, 0.5µl FW, 0.5 µL REV primer, 3µl H2O, 1µl cDNA. The reactions were run on a Bio-Rad CFX384 instrument (Bio-Rad, Hercules, CA, USA) with the following cycling parameters: initial denaturing 95°C 3min, 40 cycles of 95°C 10sec, 55°C 10sec, and 72°c 30sec with a plate read after each 72°C extension. Immediately upon PCR run, a melting curve analysis was run as follows: 95°C 10sec, 65°C-95°C with 0.5°C increments, with a plate read after each increment. The expression data were analyzed using the Bio-Rad CFX348 Maestro software with default parameters, and expression was reported as normalized expression estimated using the Comparative CT Method (ΔΔCT method) (Schmittgen and Livak, 2008).

### Data Analysis

#### Statistical Analysis and Visualization

All statistical analyses and data visualizations were conducted using R. Data wrangling and statistical modeling were performed using the dplyr and tibble packages, and visualizations were generated using the ggplot2, ggsignif, and ggpubr packages. Data transformation was done with the forecast package. Statistical analysis was done with the MASS package. ART-ANOVA was done with the ARTool package.

#### Genome-Wide Association Study (GWAS)

All statistical analyses and data visualizations were conducted using R. Data wrangling and statistical modeling were performed using the dplyr, forcats, reshape2, and tibble packages, and visualizations were generated using the ggplot2, ffpubr, corrplor, viridis, khorma, and scales packages. GWAS was performed using bigsnpr and furrr packages, and MASH was done by the mashr package.

DNA from the 138 core IPSRIs, IHP1 and ILP1 were processed by the genotype-by-sequencing (GBS) pipeline for maize (Glaubitz et al., 2014). Genotype data for the population was first filtered for minor allele frequency above 5% and a missing rate below 10% generating a set of 95,680 SNPs. Missing calls were imputed to the mean allele at that SNP and principal components (PCs) of the SNP matrix were calculated. A generalized linear model was fit for each trait in each year using the package bigsnpr in R. To establish a significance threshold, the bigsnpr function “bed_clumping()” was used to clump SNPs by LD using a threshold of R² > 0.2. This set of independent SNP’s was then used to establish a less conservative Bonferroni correction using an alpha level of .05.

After running GWAS for each trait in each year, multivariate adaptive shrinkage (mash) was run on the GWAS summary statistics. Mash is a Bayesian learning model that learns patterns of sharing among GWAS results by fitting a gaussian mixture model to effect estimates and standard errors from different traits or conditions and produces adjusted measures of significance and effect size. First, the model learns shared patterns among different GWAS results by estimating covariation between effect sizes and directions for the traits. Once this model is fit, it can be used to generate new effect estimates that are shrunk towards zero based on these learned patterns. A measure of significance for each SNP can be obtained in the form of a Bayes factor which compares the model where the SNP has zero effect vs the model where it has a non-zero effect. Bayes factors are obtained from the final model fit and used to construct a manhattan-style plot substituting p-values for Bayes factors called a mash-hattan plot. Candidate windows around mash hits were created by considering SNPs in LD with a given mash-identified SNP at R² > 0.25.

A significance threshold was established by first identifying 7,052 independent SNPs clustered by linkage disequilibrium (LD) above the threshold of R² > 0.2. 19 traits in 2014 and 23 traits in 2015 displayed ρ-values below this threshold. (Supplementary Table. 1).

To identify TAS that had statistical support across multiple years and traits, we employed a Bayesian learning model that computes posterior effect estimates and significance scores based on GWAS summary statistics (Urbut *et al*., 2019) and found 1,361 SNPs with a log10 Bayes factor > 5 (Fig. 1B). Candidate windows around mash hits were created by considering SNPs in LD with a given mash-identified SNP at R² > 0.25. Therefore, by using the IPSRI population root traits, we were able to find several root trait-associated SNPs that could contribute to the phenotypic differences in IHP1 versus ILP1 root architecture.

#### Differential expression analysis

Raw counts were used as input for DESeq2 (version X) with the full design formula: ∼ Tissue + Genotype + Treatment + Genotype:Treatment + Tissue:Genotype + Treatment:Tissue + Genotype:Treatment:Tissue. Size factors were estimated using the median ratio method and dispersions were estimated using the default DESeq2 procedure. To assess tissue-specific effects, two contrasts were extracted for root tissue. The baseline genotype effect (Fig.7 left) was estimated as the sum of the Genotype_*e1ogdh1*_vs_B73 and Tissue Root_Genotype_*e1ogdh1* coefficients, representing the genotype effect at high nitrogen specifically in root tissue. The differential nitrogen response (Fig. 7 right) was estimated as the sum of the Genotype_*e1ogdh1*_Treatment_lowN and Tissue_Root.Genotype_*e1ogdh1*.Treatment_lowN coefficients, representing the difference in nitrogen response between genotypes in root tissue. All p-values were adjusted for multiple testing using the Benjamini-Hochberg procedure. Genes with adjusted p-value < 0.05 in at least one contrast were considered differentially expressed.

#### Field Root Tip Sampling for RNA Extraction

12 plants were collected from the highest (200 bs/acre) and the lowest (50 lbs/acre) nitrogen conditions each. Root crowns were excavated and washed to expose root tips, specifically targeting aerial roots that had recently entered the soil. Approximately 2.5 cm from the distal end of living root tips was cut using sterilized razor blades and immediately transferred to 50 mL conical tubes. Each sample was dug up and cut within 10 minutes to ensure the samples reflected the natural transcriptional state. All samples were flash-frozen in liquid nitrogen in the field immediately upon collection and stored at –80°C until processing. Tissue was ground in liquid nitrogen using pre-chilled mortars and pestles before RNA extraction.

#### RNA - sequencing (RNAseq)

A total of 39 samples were processed for transcriptome sequencing, generating 261.42 Gb Clean Data. At least 6.14 Gb clean data were generated for each sample, with a minimum of 92.90% of clean data achieving a quality score of Q30. Clean reads of each sample were mapped to the specified reference genome. The mapping ratio ranged from 79.13% to 91.29%. Prediction of alternative splicing, gene structure optimization analysis, and novel gene discovery was processed on top of mapping results, during which 9,415 were discovered, and 4,540 novel genes were annotated with a putative function. Differentially expressed genes (DEGs) were identified using the criteria of Fold Change≥1.5 and FDR<0.05. For gene analysis, only genes that had an fpkm>1 for at least 2 samples were included.

## Supplementary tables

**Supplementary Table 1.**
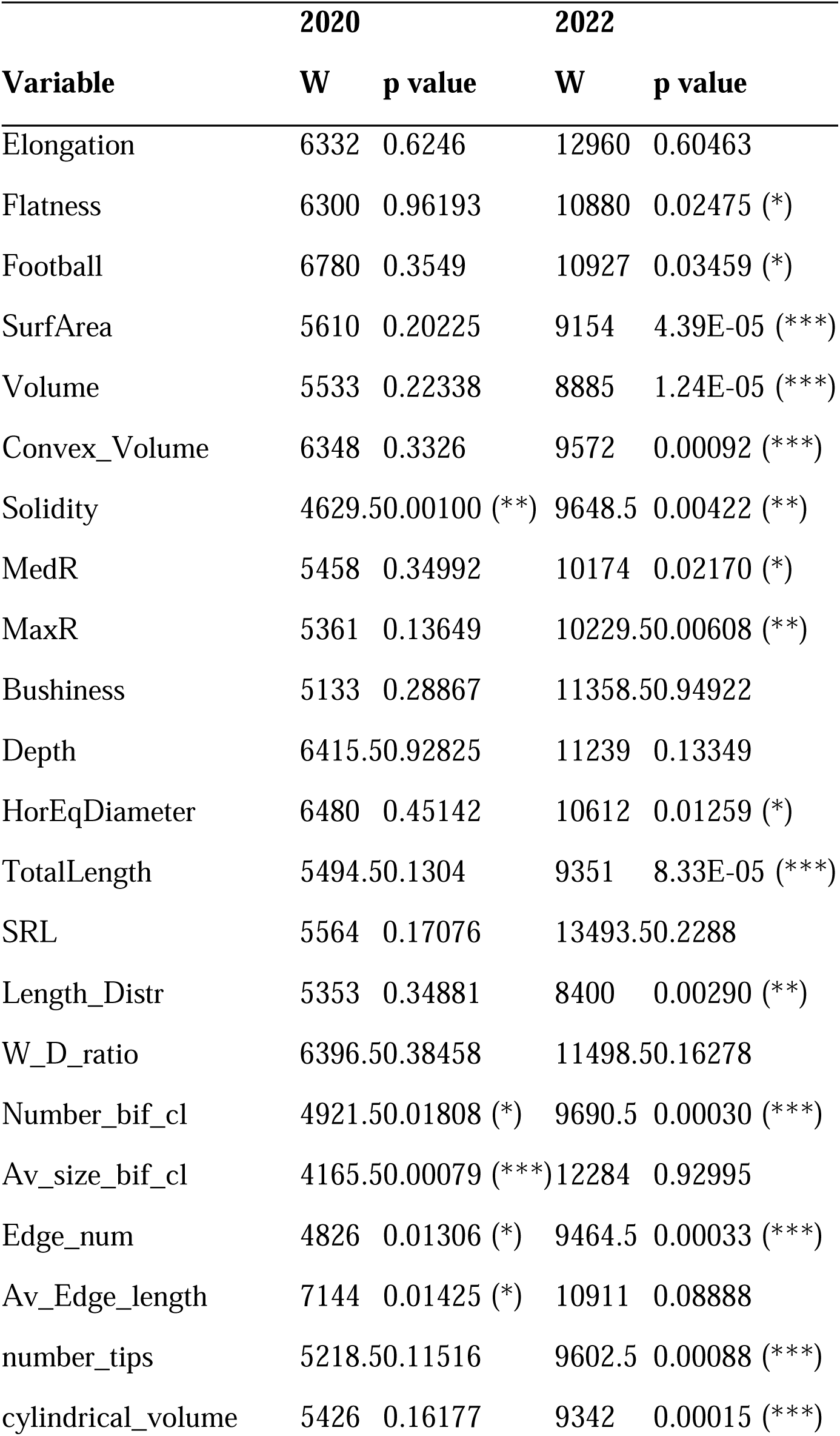
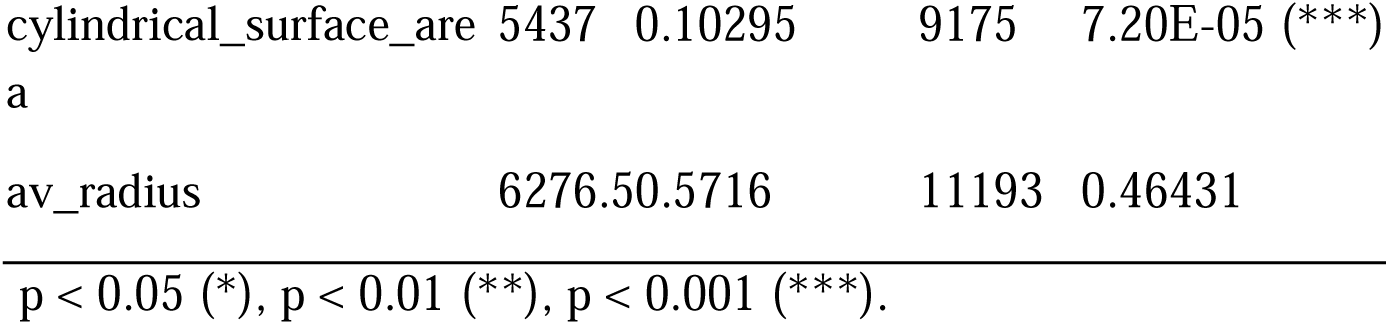
Wilcoxon p-values of 2020 and 2022 field root traits by allele.

**Supplementary Table 2.** Means of harvest index, kernel weight, nitrogen percentage, and carbon percentage for B73 and *e1ogdh*.

| Genotype | B73 | B73 | *e1ogdh1* | *e1ogdh1* |
| --- | --- | --- | --- | --- |
| N Treatment | 200 | 50 | 200 | 50 |
| Harvest index | 0.089483 | 0.025841 | 0.039486 | 0.057023 |
| 100 Kernel weight | 16.5 | 11.3 | 12.6 | 10.4 |
| N[%] | 1.11125 | 0.978571 | 1.382222 | 1.134286 |
| C[%] | 41.2425 | 40.28833 | 39.33556 | 39.16833 |

**Supplementary Table 3.** Wilcoxon test of 2024 field root traits.

|  | B73 | *e1ogdh1* | HighN | LowN |
| --- | --- | --- | --- | --- |
| Trait | High vs Low | High vs Low | B73 vs *e1ogdh1* | B73 vs *e1ogdh1* |
| Elongation | 1 | 0.0454 (*) | 0.2629 | 0.5613 |
| Flatness | 0.5959 | 0.1252 | 0.0719 | 0.1336 |
| Football | 0.5169 | 0.1832 | 0.0237 (*) | 0.0268 (*) |
| FractalDim_S | 0.0719 | 0.5635 | 0.4134 | 0.0015 (**) |
| FractalDim_T | 0.0298 (*) | 0.0104 (*) | 0.0058 (**) | 0.2243 |
| SurfArea | 0.6854 (*) | 0.0108 (*) | 0.0018 (**) | 0.0015 (**) |
| Volume | 0.3971 | 0.5254 | 0.0341 (*) | 0.0022 (**) |
| Convex_V | 0.0719 | 0.5966 | 0.1853 | 0.0240 (*) |
| Solidity | 0.1252 | 0.3253 | 0.0022 (**) | 0.0065 (**) |
| MedR | 0.5254 | 0.5241 | 0.0796 | 0.0077 (**) |
| MaxR | 0.2898 | 0.0070 (**) | 0.0004 (***) | 0.0240 (*) |
| Bushiness | 0.9485 | 0.0199 (*) | 0.3123 | 0.0538 |
| Depth | 0.2726 | 0.2678 | 0.6061 | 0.0282 (*) |
| HorEqDia | 0.0034 (**) | 0.1384 | 0.4587 | 0.0034 (**) |
| TotalLeng | 0.6025 | 0.0061 (**) | 0.0009 (***) | 0.0046 (**) |
| SRL | 0.7911 | 0.5317 | 0.8598 | 0.0371 (*) |
| Length_Di | 0.3506 | 1 | 0.665 | 0.0673 |
| W_D_ratio | 0.0034 (**) | 0.0922 | 1 | 0.0332 (*) |
| Number_bif_cl | 0.2898 | 0.0014 (**) | 0.0024 (**) | 0.0177 (*) |
| Av_size_bif_cl | 0.204 | 0.0018 (**) | 0.7911 | 0.148 |
| Edge_num | 0.2898 | 0.0014 (**) | 0.0024 (**) | 0.0728 |
| Av_Edge_length | 0.2443 | 0.0006 (***) | 0.5365 | 0.0562 |
| number_tips | 0.0566 | 0.0014 (**) | 0.0042 (**) | 0.1828 |
| Cylindrical volume | 0.3408 | 0.1182 | 0.0304 (*) | 0.0106 (*) |
| Cylindrical surface_area | 0.3253 | 0.1688 | 0.0024 (**) | 0.0022 (**) |
| av_radius | 0.9157 | 0.5244 | 0.3408 | 1 |

p < 0.05 (\*), p < 0.01 (\*\*), p < 0.001 (\*\*\*).

**Supplementary Table 4.**
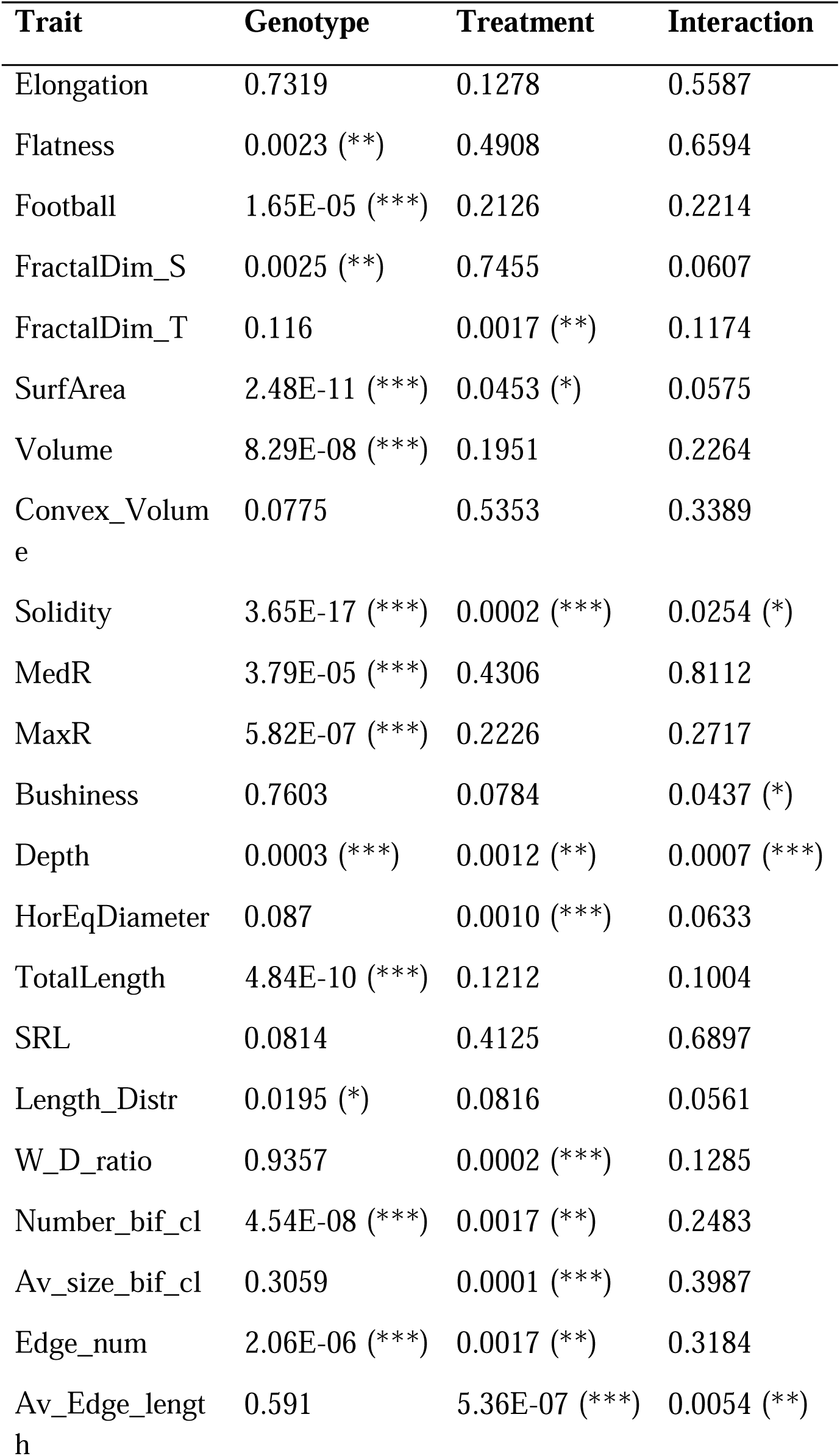
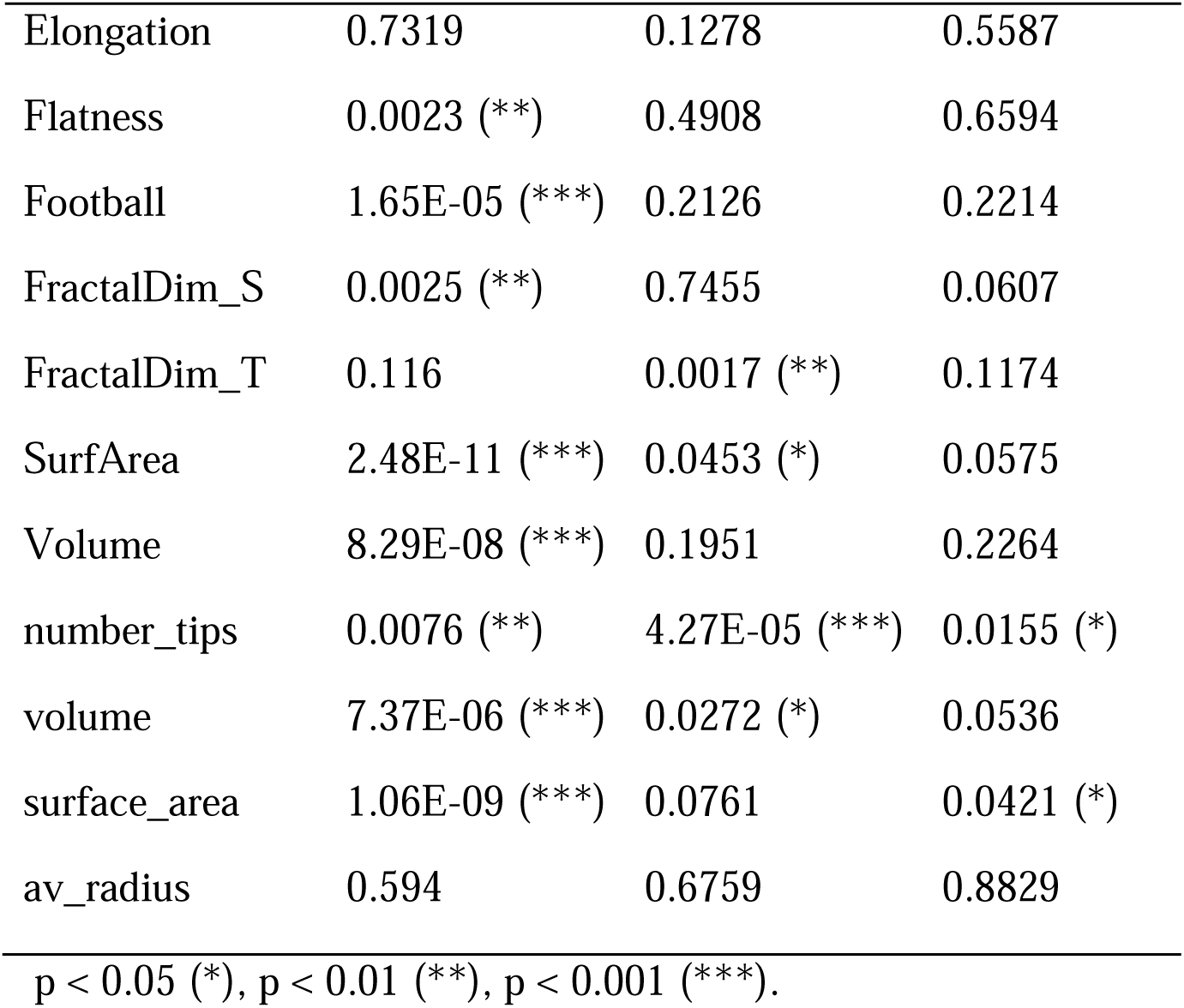
Aligned Rank Transform ANOVA (ART ANOVA) results of 2024 root traits.

## Supplementary figures

**Supplementary Figure 1:**
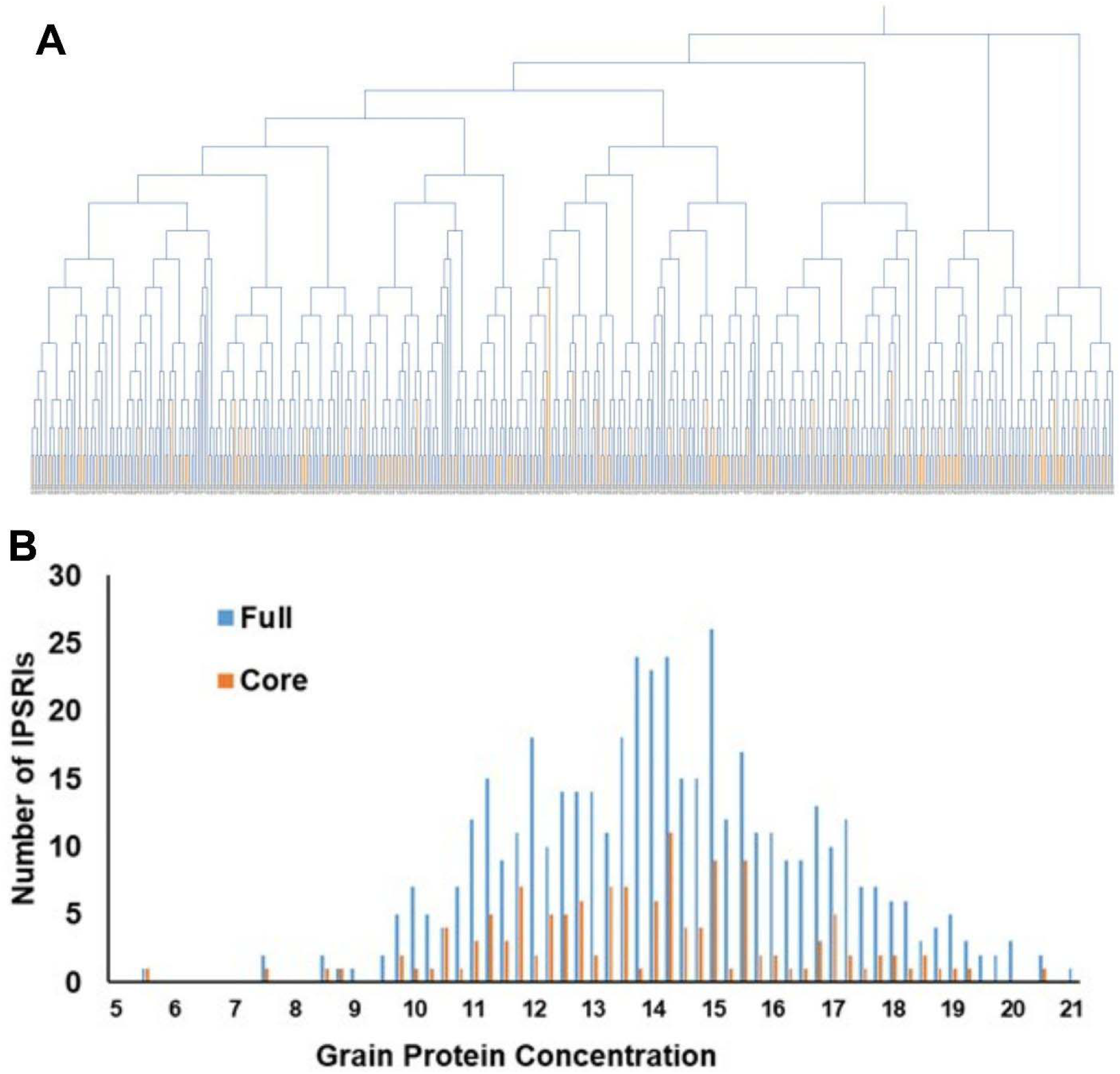
Genotypic and phenotypic diversity of the full population of 500 IPSRIs (blue) and the 138 core lines (orange). (**A**) Dendogram of IPSRIs using the 499 SNP marker genotypes from Dudley et al. (2007) and (**B**) distribution of grain protein concentration phenotype of the IPSRIs measured in 2014 field season.

**Supplementary Figure 2.**
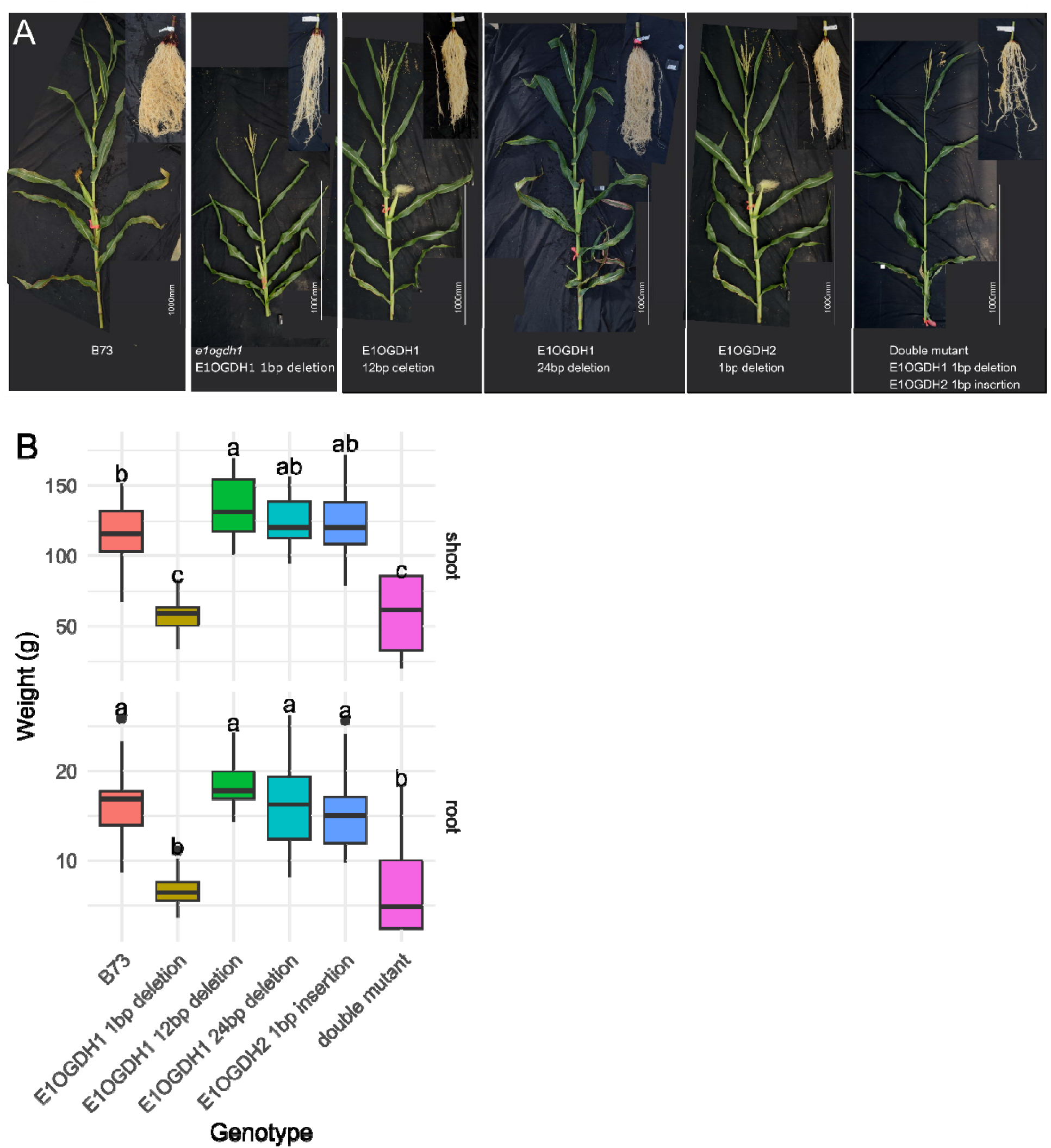
Images and biomass measurements of E1OGDH1 and E1OGDH2 CRISPR mutants shoot and root compared to B73. A. Images of B73(WT) and E1OGDH1 and E1OGDH2 mutants. The corresponding root is shown in the upper right corner of the shoot images. Images of B73 and *e1ogdh1* are identical with Figure 5B (in increased panel spaces). B. Biomass measurements of E1OGDH1 and E1OGDH2 CRISPR mutants in shoot and root. Boxplots show the distribution of shoot (top) and root (bottom) biomass (g) for B73 and CRISPR-edited lines carrying deletions or insertions in E1OGDH1 (1bp deletion, 12bp deletion, 24bp deletion) and E1OGDH2 (1bp insertion) as well as a double mutant. Different letters indicate statistically significant differences between genotypes within each tissue based on Tukey’s HSD test (p < 0.05). Center line represents the median, box boundaries represent the interquartile range, and whiskers extend to 1.5× the interquartile range.

## Author contributions

M.C. designed and conducted the 2020, 2022, and 2024 field experiments, performed data analysis and statistical interpretation, and drafted the manuscript, with inputs from C.N.T. and S.M. M.C. interpreted the results with input from C.N.T. Z.L. and C.N.T. conceived the project and collected samples from the 2014 and 2015 field experiments. S.M. and C.T. provided samples from the 2014 and 2015 field, and plant materials. Z.L. performed the initial genome-wide association study (GWAS), with assistance of C.L., and G.Z. C.L. and I.B. performed further multi-year GWAS. K.T. provided genotyping-by-sequencing (GBS) data for the analysis. M.C. and D.T. conducted molecular biology experiments. B.Y. and H.L. generated and provided the CRISPR/Cas9 mutants. All authors reviewed and approved the final manuscript.

## Acknowledgments

The authors would like to thank Catherine Li (Moose lab) for helping make the IPSRI supplementary figure and Alexandria Tran (Moose lab) for help with the E1OGDH1 sequencing analysis. They would also like to thank the Moose, Baxter, and Topp field teams for their tireless contributions to data collection.

## Funding

This material is based upon work supported by the National Science Foundation under Award number IOS-1638507 and by the the Donald Danforth Plant Science Center’s Subterranean

Influences on Nitrogen and Carbon Cycling (SINC) Center.

## Works Cited

Alrefai, R., Orozco, B. and Rocheford, T. (1994) “Detection and sequencing of the transposable element ILS-1 in the Illinois long-term selection maize strains,” Plant physiology, 106(2), pp. 803–804.

Baer, M. et al. (2023) “Maize lateral rootless 1 encodes a homolog of the DCAF protein subunit of the CUL4-based E3 ubiquitin ligase complex,” The New Phytologist, 237(4), pp. 1204–1214.

Bray, A.L. and Topp, C.N. (2018) “The Quantitative Genetic Control of Root Architecture in Maize,” Plant & cell physiology, 59(10), pp. 1919–1930.

Char, S.N. et al. (2017) “An Agrobacterium-delivered CRISPR/Cas9 system for high-frequency targeted mutagenesis in maize,” Plant biotechnology journal, 15(2), pp. 257–268.

Cheng, Y. et al. (2018) “Glutamate receptor Homolog3.4 is involved in regulation of seed germination under salt stress in Arabidopsis,” Plant & cell physiology, 59(5), pp. 978–988.

Condori-Apfata, J.A. et al. (2019) “The Arabidopsis E1 subunit of the 2-oxoglutarate dehydrogenase complex modulates plant growth and seed production,” Plant molecular biology, 101(1–2), pp. 183–202.

Das, A. et al. (2015) “Digital imaging of root traits (DIRT): a high-throughput computing and collaboration platform for field-based root phenomics,” Plant methods, 11, p. 51.

Dechorgnat, J. et al. (2019) “Tissue and nitrogen-linked expression profiles of ammonium and nitrate transporters in maize,” BMC plant biology, 19(1), p. 206.

Dudley, J.W. (2007) “From means to QTL: The Illinois long-term selection experiment as a case study in quantitative genetics,” Crop science, 47(Supplement_3), p. S-20–S-31.

Feng, Z. et al. (2025) “ZmGCT1/2 negatively regulate drought tolerance in maize by inhibiting ZmSLAC1 to maintain guard cell turgor,” Proceedings of the National Academy of Sciences of the United States of America, 122(15), p. e2423037122.

Forde, B.G. (2002) “Local and long-range signaling pathways regulating plant responses to nitrate,” Annual review of plant biology, 53(1), pp. 203–224.

Forde, B.G. et al. (2013) “Glutamate signalling via a MEKK1 kinase-dependent pathway induces changes in Arabidopsis root architecture,” The Plant journal: for cell and molecular biology, 75(1), pp. 1–10.

Forde, B.G. and Walch-Liu, P. (2009) “Nitrate and glutamate as environmental cues for behavioural responses in plant roots,” Plant, cell & environment, 32(6), pp. 682–693.

Frank, R.A.W. et al. (2007) “Crystal structure of the E1 component of the Escherichia coli 2-oxoglutarate dehydrogenase multienzyme complex,” Journal of molecular biology, 368(3), pp. 639–651.

Gallais, A. and Hirel, B. (2004) “An approach to the genetics of nitrogen use efficiency in maize,” Journal of experimental botany, 55(396), pp. 295–306.

Gao, K. et al. (2015) “A comprehensive analysis of root morphological changes and nitrogen allocation in maize in response to low nitrogen stress,” Plant, cell & environment, 38(4), pp. 740–750.

Gaudin, A.C.M. et al. (2011) “Novel temporal, fine-scale and growth variation phenotypes in roots of adult-stage maize (Zea mays L.) in response to low nitrogen stress: Nitrogen stress on maize roots,” Plant, cell & environment, 34(12), pp. 2122–2137.

Gaudin, A.C.M., McClymont, S.A. and Raizada, M.N. (2011) “The nitrogen adaptation strategy of the wild teosinte ancestor of modern maize,Zea mayssubsp.Parviglumis,” Crop science, 51(6), pp. 2780–2795.

Goldman, I.L., Rocheford, T.R. and Dudley, J.W. (1993) “Quantitative trait loci influencing protein and starch concentration in the Illinois Long Term Selection maize strains,” TAG. Theoretical and applied genetics. Theoretische und angewandte Genetik, 87(1–2), pp. 217–224.

Goldman, I.L., Rocheford, T.R. and Dudley, J.W. (1994) “Molecular markers associated with maize kernel oil concentration in an Illinois high protein× Illinois low protein cross,” Crop science [Preprint]. Available at: https://acsess.onlinelibrary.wiley.com/doi/abs/10.2135/cropsci1994.0011183X003400040013x.

Good, A.G. and Beatty, P.H. (2011) “Fertilizing nature: a tragedy of excess in the commons,” PLoS biology, 9(8), p. e1001124.

Haelterman, L. et al. (2025) “Genetic control of root morphology in rapeseed recombinant inbred lines grown under contrasting nitrogen levels,” Physiologia plantarum, 177(4), p. e70431.

Han, B. et al. (2012) “ZmHO-1, a maize haem oxygenase-1 gene, plays a role in determining lateral root development,” Plant Science: An International Journal of Experimental Plant Biology, 184, pp. 63–74.

Hein, K.M. et al. (2025) “Phenome-to-genome insights for evaluating root system architecture in field studies of maize,” The Plant Genome, 18(3), p. e70100.

Hoener, I.R. and DeTurk, E.E. (1938) “The absorption and utilization of nitrate nitrogen during vegetative growth by Illinois high protein and Illinois low protein corn 1,” Agronomy journal, 30(3), pp. 232–243.

Jia, L. et al. (2023) “Genome-wide identification and functional analysis of nitrate transporter genes (NPF, NRT2 and NRT3) in maize,” International journal of molecular sciences, 24(16), p. 12941.

Kan, C.-C. et al. (2017) “Exogenous glutamate rapidly induces the expression of genes involved in metabolism and defense responses in rice roots,” BMC genomics, 18(1), p. 186.

Kirschner, G.K. et al. (2024) “Genetic regulation of the root angle in cereals,” Trends in Plant Science, 29(7), pp. 814–822.

Krapp, A. et al. (2011) “Arabidopsis roots and shoots show distinct temporal adaptation patterns toward nitrogen starvation,” Plant physiology, 157(3), pp. 1255–1282.

Li, D. et al. (2025) “Maize ZmDLR2/BRU1 is required for lateral root primordium emergence by participating in DNA repair,” *Plant*, Cell & Environment, 48(10), pp. 7377–7391.

Li, H. et al. (2009) “The Sequence Alignment/Map format and SAMtools,” *Bioinformatics (Oxford*, England), 25(16), pp. 2078–2079.

Li, H. (2018) “Minimap2: pairwise alignment for nucleotide sequences,” *Bioinformatics (Oxford*, England), 34(18), pp. 3094–3100.

Li, M. et al. (2018) “The persistent homology mathematical framework provides enhanced genotype-to-phenotype associations for plant morphology,” Plant Physiology, 177(4), pp. 1382– 1395.

Li, M. et al. (2023) “Topological data analysis expands the genotype to phenotype map for 3D maize root system architecture,” Frontiers in plant science, 14, p. 1260005.

Li, P. et al. (2019) “Natural variation of ZmHKT1 affects root morphology in maize at the seedling stage,” Planta, 249(3), pp. 879–889.

Li, S. et al. (2024) “NRG2 family members of Arabidopsis and maize regulate nitrate signalling and promote nitrogen use efficiency,” Physiologia plantarum, 176(2), p. e14251.

Li, Z. et al. (2018) “Enhancing auxin accumulation in maize root tips improves root growth and dwarfs plant height,” Plant Biotechnology Journal, 16(1), pp. 86–99.

Li, Z.-G., Ye, X.-Y. and Qiu, X.-M. (2019) “Glutamate signaling enhances the heat tolerance of maize seedlings by plant glutamate receptor-like channels-mediated calcium signaling,” Protoplasma, 256(4), pp. 1165–1169.

Liu, L. et al. (2023) “Varietal responses of root characteristics to low nitrogen application explain the differing nitrogen uptake and grain yield in two rice varieties,” Frontiers in plant science, 14, p. 1244281.

Liu, S. et al. (2021) “DIRT/3D: 3D root phenotyping for field-grown maize (Zea mays),” Plant Physiology, 187(2), pp. 739–757.

Lohaus, G. et al. (1998) “Transport of amino acids with special emphasis on the synthesis and transport of asparagine in the Illinois Low Protein and Illinois High Protein strains of maize,” Planta, 205(2), pp. 181–188.

Lucas, C.J. et al. (2013) “Genomic changes in response to 110 cycles of selection for seed protein and oil concentration in maize,” in Seed Genomics. Oxford, UK: Wiley-Blackwell, pp. 217–236.

Ma, H. et al. (2022) “Poly-γ-glutamic acid promoted maize root development by affecting auxin signaling pathway and the abundance and diversity of rhizosphere microbial community,” BMC plant biology, 22(1), p. 521.

Ma, N. et al. (2020) “Transcriptome analysis of maize seedling roots in response to nitrogen-, phosphorus-, and potassium deficiency,” Plant and Soil, 447(1–2), pp. 637–658.

Mann, C. (1997) “Reseeding the Green revolution,” Science (New York, N.Y.), 277(5329), pp. 1038–1043.

Masclaux-Daubresse, C. et al. (2006) “Glutamine synthetase-glutamate synthase pathway and glutamate dehydrogenase play distinct roles in the sink-source nitrogen cycle in tobacco,” Plant physiology, 140(2), pp. 444–456.

Messina, C. et al. (2021) “Reproductive resilience but not root architecture underpins yield improvement under drought in maize,” Journal of experimental botany, 72(14), pp. 5235–5245.

Mikkilineni, V. and Rocheford, T.R. (2004) “RFLP variant frequency differences among Illinois long-term selection protein strains,” Plant breeding reviews, 24(1), pp. 111–132.

Millar, A.H., Hill, S.A. and Leaver, C.J. (1999) “Plant mitochondrial 2-oxoglutarate dehydrogenase complex: purification and characterization in potato,” The Biochemical journal, 343 Pt 2, pp. 327–334.

Miller, A.J. et al. (2008) “Amino acids and nitrate as signals for the regulation of nitrogen acquisition,” Journal of experimental botany, 59(1), pp. 111–119.

Moose, S.P., Dudley, J.W. and Rocheford, T.R. (2004) “Maize selection passes the century mark: a unique resource for 21st century genomics,” Trends in plant science, 9(7), pp. 358–364.

Mueller, S.M., Messina, C.D. and Vyn, T.J. (2019) “Simultaneous gains in grain yield and nitrogen efficiency over 70 years of maize genetic improvement,” Scientific reports, 9(1), p. 9095.

Nedelyaeva, O.I. et al. (2024) “Functional and molecular characterization of plant nitrate transporters belonging to NPF (NRT1/PTR) 6 subfamily,” International Journal of Molecular Sciences, 25(24), p. 13648.

Philippe, F. et al. (2019) “Involvement of Medicago truncatula glutamate receptor-like channels in nitric oxide production under short-term water deficit stress,” Journal of plant physiology, 236, pp. 1–6.

Postma, J.A., Dathe, A. and Lynch, J.P. (2014) “The optimal lateral root branching density for maize depends on nitrogen and phosphorus availability,” Plant physiology, 166(2), pp. 590–602.

Pradhan, P. et al. (2015) “Closing yield gaps: How sustainable can we be?,” PloS one, 10(6), p. e0129487.

Qi, Z., Stephens, N.R. and Spalding, E.P. (2006) “Calcium entry mediated by GLR3.3, an Arabidopsis glutamate receptor with a broad agonist profile,” Plant physiology, 142(3), pp. 963– 971.

Qin, L. et al. (2025) “Phenotypic characterization and transcriptome analysis of the dwarf mutant zmbrd1 in maize,” Genes, 16(12), p. 1410.

Raun, W.R. and Johnson, G.V. (1999) “Improving nitrogen use efficiency for cereal production,” Agronomy journal, 91(3), pp. 357–363.

Ren, W. et al. (2022) “Genome-wide dissection of changes in maize root system architecture during modern breeding,” Nature Plants, 8(12), pp. 1408–1422.

Ruiz, A. et al. (2023) “Harvest index has increased over the last 50 years of maize breeding,” Field crops research, 300(108991), p. 108991.

Schmittgen, T.D. and Livak, K.J. (2008) “Analyzing real-time PCR data by the comparative C(T) method,” Nature protocols, 3(6), pp. 1101–1108.

Schneider, H.M. et al. (2021) “Nodal root diameter and node number in maize (Zea mays L.) interact to influence plant growth under nitrogen stress,” Plant direct, 5(3), p. e00310.

Sciarresi, C. et al. (2025) “Breeding for high maize yields indirectly boosting root carbon in the US Corn Belt since the 1980s,” Field crops research, 323(109774), p. 109774.

Shao, M.R. et al. (2021) “Complementary Phenotyping of Maize Root System Architecture by Root Pulling Force and X-Ray Imaging,” Plant phenomics (Washington, D.C.), 2021, p. 9859254.

Singh, T. et al. (2024) “Paenibacillus lentimorbus alleviates nutrient deficiency-induced stress in Zea mays by modulating root system architecture, auxin signaling, and metabolic pathways,” Plant Cell Reports, 43(2), p. 49.

Suzuki, M. et al. (2015) “Conserved functions of the MATE transporter BIG EMBRYO1 in regulation of lateral organ size and initiation rate,” The Plant Cell, 27(8), pp. 2288–2300.

Tilman, D. et al. (2002) “Agricultural sustainability and intensive production practices,” Nature, 418(6898), pp. 671–677.

Topp, C.N. et al. (2016) “How can we harness quantitative genetic variation in crop root systems for agricultural improvement?: Quantifying root architecture for crops,” Journal of Integrative Plant Biology, 58(3), pp. 213–225.

Trachsel, S. et al. (2013) “Maize root growth angles become steeper under low N conditions,” Field crops research, 140, pp. 18–31.

Urbut, S.M. et al. (2019) “Flexible statistical methods for estimating and testing effects in genomic studies with multiple conditions,” Nature genetics, 51(1), pp. 187–195.

Uribelarrea, M., Below, F.E. and Moose, S.P. (2004) “Grain composition and productivity of maize hybrids derived from the Illinois protein strains in response to variable nitrogen supply,” Crop science, 44(5), pp. 1593–1600.

Uribelarrea, M., Crafts-Brandner, S.J. and Below, F.E. (2008) “Physiological N response of field-grown maize hybrids (Zea mays L.) with divergent yield potential and grain protein concentration,” Plant and soil, 316(1), p. 151.

Vincill, E.D., Bieck, A.M. and Spalding, E.P. (2012) “Ca(2+) conduction by an amino acid-gated ion channel related to glutamate receptors,” Plant physiology, 159(1), pp. 40–46.

Walch-Liu, P. et al. (2006) “Evidence that L-glutamate can act as an exogenous signal to modulate root growth and branching in Arabidopsis thaliana,” Plant & cell physiology, 47(8), pp. 1045–1057.

Wei, N. et al. (2023) “Characterization of the isocitrate dehydrogenase gene family and their response to drought stress in maize,” Plants, 12(19), p. 3466.

West, P.C. et al. (2014) “Leverage points for improving global food security and the environment,” Science (New York, N.Y.), 345(6194), pp. 325–328.

York, L.M. and Lynch, J.P. (2015) “Intensive field phenotyping of maize (Zea mays L.) root crowns identifies phenes and phene integration associated with plant growth and nitrogen acquisition,” Journal of experimental botany, 66(18), pp. 5493–5505.

Zhang, C. et al. (2026) “The maize ZmbHLH118 transcription factor regulates vacuolar nitrate loading by the NO3 - transporter ZmCLCa,” Advanced Science (Weinheim, Baden-Wurttemberg, Germany), 13(25), p. e20219.

Zhang, X. et al. (2020) “Genetic variation in ZmTIP1 contributes to root hair elongation and drought tolerance in maize,” Plant biotechnology journal, 18(5), pp. 1271–1283.

Zhang, Y. et al. (2022) “Morphological characterization and transcriptome analysis of leaf angle mutant bhlh112 in maize [Zea mays L.],” Frontiers in Plant Science, 13, p. 995815.

Zhang, Z., Chen, L. and Yu, J. (2023) “Maize WRKY28 interacts with the DELLA protein D8 to affect skotomorphogenesis and participates in the regulation of shade avoidance and plant architecture,” Journal of Experimental Botany, 74(10), pp. 3122–3141.

Zheng, Y. et al. (2018) “The glutamate receptors AtGLR1.2 and AtGLR1.3 increase cold tolerance by regulating jasmonate signaling in Arabidopsis thaliana,” Biochemical and biophysical research communications, 506(4), pp. 895–900.

Zhu, Z.L. and Chen, D.L. (2002) “Nitrogen fertilizer use in China – Contributions to food production, impacts on the environment and best management strategies,” Nutrient cycling in agroecosystems, 63(2/3), pp. 117–127.

